# Functional convergence in slow-growing microbial communities arises from thermodynamic constraints

**DOI:** 10.1101/2022.03.11.483989

**Authors:** Ashish B. George, Tong Wang, Sergei Maslov

## Abstract

The dynamics of microbial communities is complex, determined by competition for metabolic substrates and cross-feeding of byproducts. Species in the community grow by harvesting energy from chemical reactions that transform substrates to products. In many anoxic environments, these reactions are close to thermodynamic equilibrium and growth is slow. To understand the community structure in these energy-limited environments, we developed a microbial community consumer-resource model incorporating energetic and thermodynamic constraints on an interconnected metabolic network. The central element of the model is product inhibition, meaning that microbial growth may be limited not only by depletion of metabolic substrates but also by accumulation of products. We demonstrate that these additional constraints on microbial growth cause a convergence in the structure and function of the community metabolic network—independent of species composition and biochemical details—providing a possible explanation for convergence of community function despite taxonomic variation observed in many natural and industrial environments. Furthermore, we discovered that the structure of community metabolic network is governed by the thermodynamic principle of maximum free energy dissipation. Our results predict the decrease of functional convergence in faster growing communities, which we validate by analyzing experimental data from anaerobic digesters. Overall, the work demonstrates how universal thermodynamic principles may constrain community metabolism and explain observed functional convergence in microbial communities.

## INTRODUCTION

Over half of Earth’s prokaryotic biomass resides in ocean sediments and deep soil^1,2^.In many such natural environments and industrial anaerobic digesters, microbial communities grow under energy-limited conditions due to the limited availability of strong electron acceptors, like oxygen or nitrate, and strong electron donors, like glucose^3–9^. Hence species in these communities are forced to harvest energy required for biomass growth from low free energy reactions, such as anaerobic respiration using sulphate or carbon dioxide and certain fermentation reactions^9–14^. Growth in these environments can be very slow—division times are measured in weeks and years—instead of minutes and hours used for growth in the lab^4, 14–17^. These long timescales and difficulty in culturing these species in a lab environment make experimental investigation of these communities challenging. Models and theory of energy-limited microbial communities can help uncover their organizational principles and shared emergent properties and drive understanding.

Models and theories inspired by MacArthur’s consumer-resource model have helped understand many of the principles and properties governing microbial communities structured by nutrient-limitation^18–24^. However, models of communities in low-energy environments, structured by energy-limitation, need to incorporate important additional details. First, microbial growth in low-energy environments is determined by energy assimilation rather than nutrient uptake ^25–27^. Second, the low free energy reactions utilized to drive growth are subject to thermodynamic constraints. And third, reactions need to be explicitly modeled to track their thermodynamic feasibility, rather than just species preference for resources. Here we propose and study a minimal model of microbial communities that accounts for these crucial details to understand the emergent properties of slow-growing communities in energy-limited environments.

Recently, several studies have attempted to incorporate thermodynamics into consumer-resource style microbial community models^28, 29^. These studies focused on community diversity, showing that thermodynamics enables communities to overcome the competitive exclusion principle and stabilize high diversity on a single substrate. In contrast, here we focus on the emergent properties of the communities in low-energy environments, such as the metabolic network structure and function of the community.

Simulations of our model generated communities that shared many features of natural communities. Notably, communities assembled from separate species pools in replicate environments realized the same metabolic network and performed similar metabolic functions at the community-level despite stark differences at the species level. This resembles the observed convergence in community function despite taxonomic divergence, seen in anaerobic bioreactors, oceans, and other environments^30–33^.

Analyzing the model to understand how functional convergence arises, we discovered that thermodynamic principles govern community-level metabolic network structure and function in slow-growing, energy-limited communities. We derived the thermodynamic principle of maximum free energy dissipation, which, we show, determines the community metabolic network selected by ecological competition in energy-limited communities. This thermodynamic principle, along with physical constraints, shape the metabolic environment created by the community and the metabolic functions it performs. Further, the derivation of the maximum dissipation principle provides a concrete example in which communities are structured by thermodynamic optimization, an idea that has been conjectured in ecology but never proven^34^.

Overall, our results highlight the crucial role of thermodynamic constraints in defining community metabolism and function of energy-limited microbial communities. The results provide an explanation for the observed convergence in microbial community function, serve as a concrete example of thermodynamic optimization in community ecology, and make predictions for the community metabolic network and resource environment.

## RESULTS

### Model of microbial communities in energy-poor environments

To understand slow-growing microbial communities in energy-limited environments, we develop a model that incorporates crucial features of growth in these environments: energy-limited growth, thermodynamic constraints on reactions, and explicit modeling of reaction fluxes. In the model, species grow utilizing energy harvested from chemical reactions that convert a higher energy resource (substrate) to a lower energy resource (product) (Fig. 1A). A reaction can be understood as a coarse-grained description of catabolism^35, 36^. The resources consumed or produced by the community are labeled in the order of decreasing standard state energy as *R*_0_*, R*_1_*, R*_2_*,…*. A species can utilize reactions connecting any pair of these resources; the allowed reactions make a directed, fully connected network. The product of one reaction can act as the substrate of another, facilitating cross-feeding interactions between microbes.

**FIG. 1.**
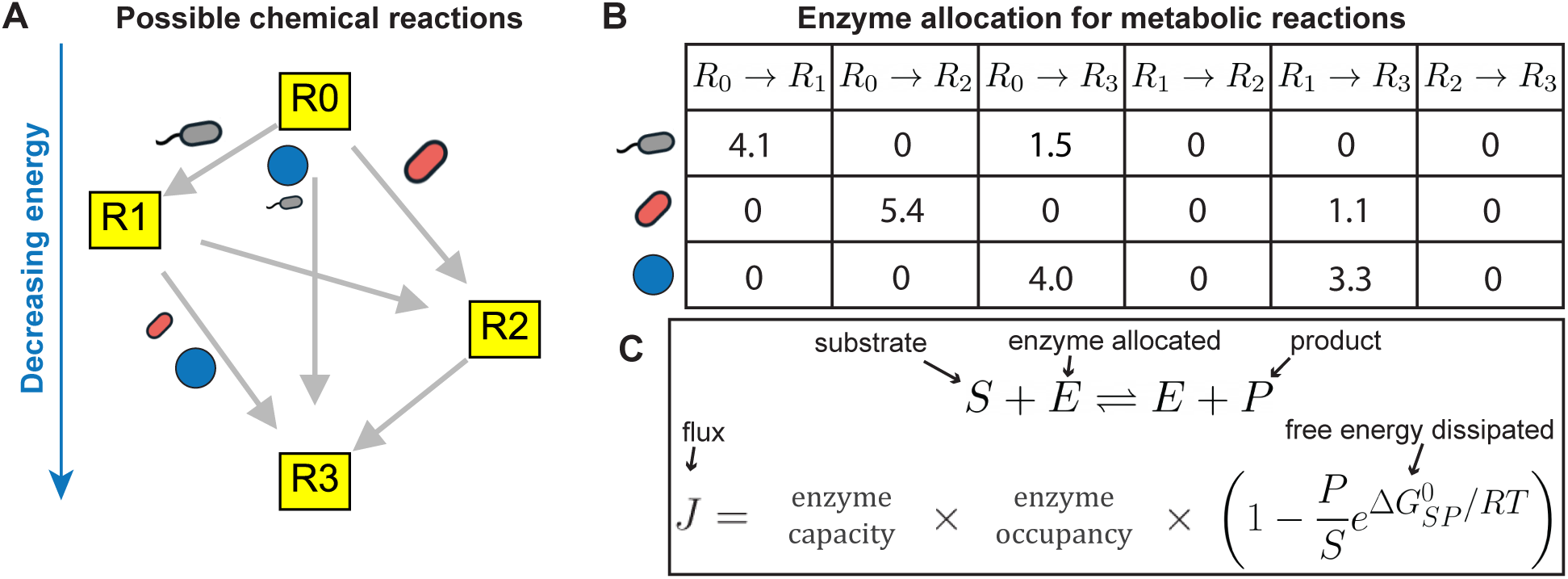
Model of communities in energy-limited environments. **A)** Species harvest energy from reactions (gray arrows) that consume a higher energy resource (substrate) to produce a lower energy resource (product). Reactions connecting any pair of the four resources *R*_0_*, R*_1_*, R*_2_*, R*_3_ can be utilized by a species, making the space of all possible reactions a fully connected network. **B)** A species can utilize a reaction by allocating a part of its overall enzyme budget to the corresponding, reaction-specific enzyme. Species differ in both the manner of enzyme allocation and their overall enzyme budgets. **C)** The reactions in low-energy environments are thermodynamically reversible. The flux *J* in a reversible, enzyme-catalyzed reaction between substrate *S* and product *P* is determined by three factors: 1) Enzyme capacity, which is the amount of enzyme available. 2) Enzyme occupancy, which measure the fraction of free enzyme available to catalyze the reaction, determined by the concentration of substrates and products. 3) Thermodynamic inhibition due to accumulation of products. Since low-energy reactions are reversible, the net flux in the forward direction will decrease as products accumulate. The strength of this inhibition is controlled by the free energy dissipated in the reaction, *−*Δ*G*^0^.

Species utilize a reaction by producing the corresponding, reaction-specific enzyme. A species allocates a portion of its total enzyme budget to any number of possible reactions. We allow total enzyme budgets to vary, as in real species^37^; this also avoids potential degenerate behavior seen in community models with fixed enzyme budgets^19^. Thus, a species is characterized by its enzyme allocation strategy and total enzyme budget (Fig. 1B). Through the explicit connection of species to reactions, substrate consumption can create different products depending on the enzyme used, like in real microbes^38^. This contrasts with microbial consumer-resource models, where substrate consumption by a species leads to a fixed mixture of products, i.e., all species use the same reactions and differ only in their consumption preferences^22, 39, 40^.

From a reaction converting substrate *S* to product *P* with standard-state energies 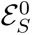 and 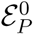 a species assimilates 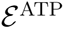 of energy. This energy is stored as ATP or in other forms of energy-rich intermediates ^41^. Since such energy assimilation will be constrained by reaction stoichiometry and biochemistry, we assume 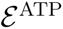 to depend only on the particular reaction and independent of the species.

The harvested energy drives the proportional growth of new biomass. The remaining free energy, 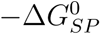, is dissipated. 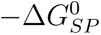 is defined as

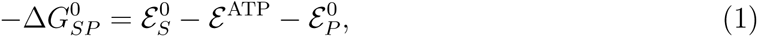

The flux through the reaction is determined by reversible Michaelis-Menten kinetics^42^:

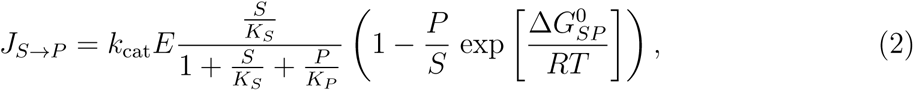

where *E* is the enzyme concentration, *k*_cat_*, K_S_, K_P_* are enzyme-specific parameters describing the chemical kinetics, *T* is the temperature, and *R* is the gas constant. In thermodynamic parlance, 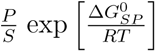 is the Boltzmann factor of the free energy difference of the reaction after accounting for resource concentration gradient, 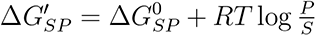. Note that our results will apply to a general stoichiometric ratio of substrate and product beyond 1: 1 as well (see SI); we omit discussion of this scenario in the main text for clarity.

Eq. 2 can be understood as a product of three terms^43^. The first term specifies the maximal reaction rate, which is proportional to the amount of enzyme allocated to the reaction, *E*. The second term describes the saturation level of the enzyme; the flux increases linearly at low substrate concentrations and saturates at high substrate concentrations. The last (and most crucial for our purposes) term describes thermodynamic inhibition, i.e., how the net rate of a reversible reaction decreases as products accumulate and the ratio *P/S* increases. The strength of the thermodynamic inhibition is controlled by the free energy dissipated in the reaction. This thermodynamic inhibition of microbial growth has been quantitatively validated in the lab and in situ conditions^6, 7, 44–47^. If product accumulation is severe enough to reverse the reaction direction, we assume that species stop enzyme production to prevent energy loss from catalyzing the reverse reaction.

The dynamics of species and resources are driven by these reaction fluxes. Species grow in proportion to the energy they assimilate from the reactions catalyzed by their enzymes. The concentration of a resource decreases due to consumption in a reaction and increases due to production in a reaction. Additionally, the entire system is diluted at a rate *δ*, which represents the dilution rate of the bioreactor/chemostat or the dilution by sedimentation in oceans^48^. Mathematically, the term representing dilution of species biomass can also be defined to account for the maintenance energy requirements of a species (see SI Sec. S5). The precise equations describing species and resource dynamics are detailed in Methods. We also obtained similar results in a model without explicit dilution of resources that incorporates species maintenance costs (see SI Sec. S7); we omit discussion of this model in the main text for clarity.

Communities in natural and industrial environments are the result of years of ecological competition and succession. To model such communities that emerge from ecological competition, we studied community assembly from a large and diverse species pool in an environment consisting of 6 resources, labelled in the order of decreasing standard-state energies as *R*_0_*, R*_1_*…, R*_5_. The environment was supplied with the most energy-rich resource *R*_0_ at high concentrations compared to the typical *K_S_*. The other lower energy resources were generated as products in some of the 15 reactions that could be utilized by species for growth. The energy assimilated from a reaction was chosen to be a random fraction of the energy gap between product and substrate of that reaction (see Methods). Enzymatic parameters describing reaction kinetics, were chosen from lognormal distributions, motivated by empirical observations^49^. Since we are interested in slow-growing communities, we chose a small dilution rate so as to not drive any slow-growing species to extinction. After introducing species from the pool into the environment, many species went extinct from competition and simulations settled down to a steady-state community comprised of the survivors.

### Emergent metabolic structure and function in slow-growing, energy-limited com-munities

To study the features shared across slow-growing communities in energy-limited environments, we repeated the community assembly experiment from five separate pools of 600 species each. Each species is characterized by its enzyme budget, chosen from a lognormal distribution, fractionally allocated to a random subset of 15 possible reactions. Species were not shared across pools. We also varied enzymatic parameters (chemical kinetic parameters) to vary between pools. The environment and its energetics was shared across the experiments, i.e., the resource supply, resource energies, and energy assimilated in each reaction (which is constrained by stoichiometry) was the same across experiments. Simulations of all pools converged to a steady-state maintained by a community of the surviving species. This resembles observations in many microbial community models and experiments^50^.

We analyzed the final steady-state communities to find emergent properties of energy-limited communities that were shared across pools that differed in species and enzyme content. We identified three properties of the communities that resemble observations in natural communities:

#### Concurrent community metabolic networks

Only a subset of the 15 possible reactions were active (i.e., carried nonzero flux) in the final communities (colored arrows in Fig. 2A). This subset of reactions was shared across communities, despite starting from different species pools (Fig. 2A,B). Hence, the reactions utilized by the surviving species were determined by the environment. Furthermore, the metabolic network structure was constrained. The in-degree of each node was 1, i.e., each resource was produced by only a single reaction.

**FIG. 2.**
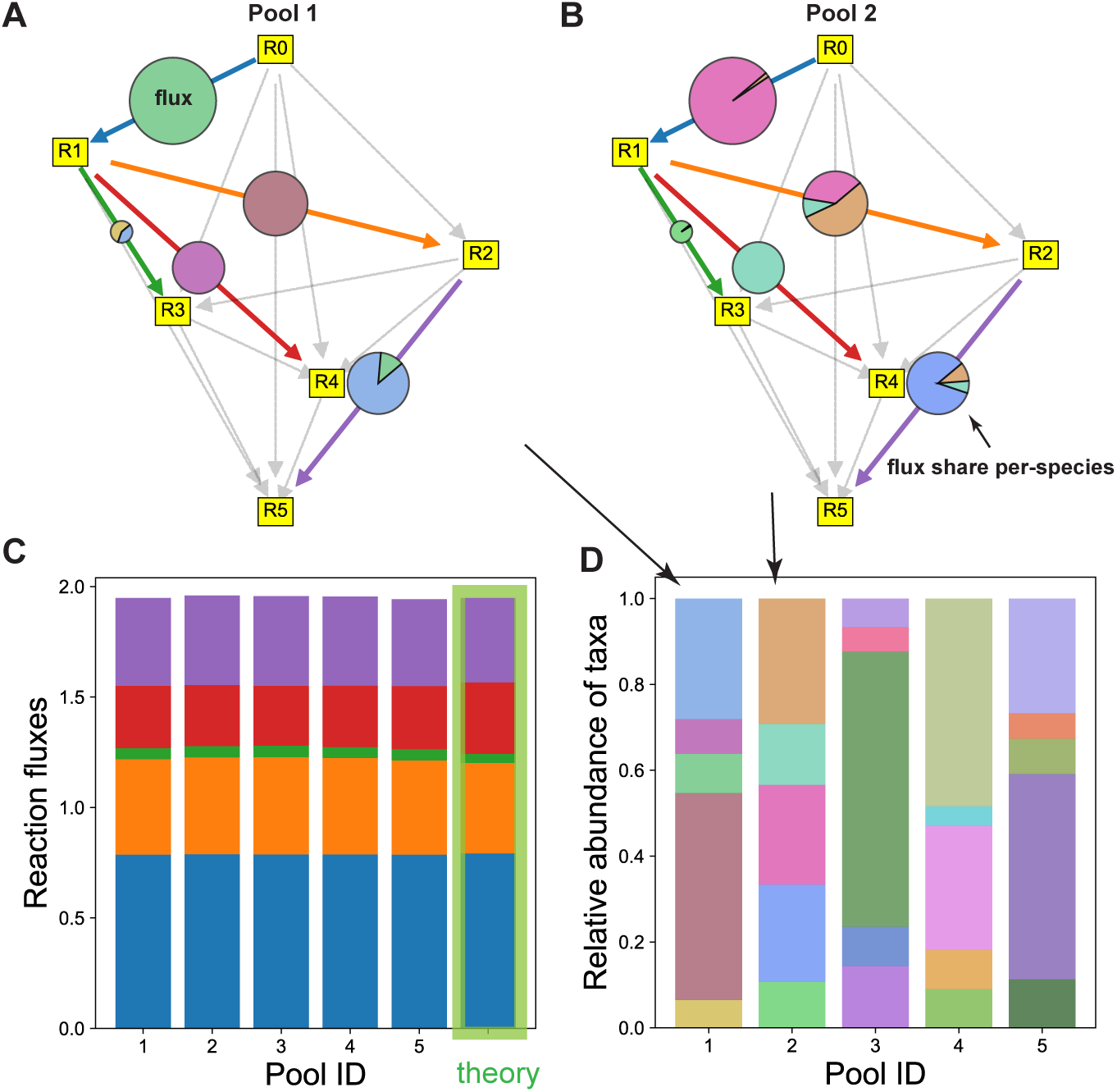
Functional convergence despite taxonomic variation. We simulated the model to obtain the final community that arose as the outcome of competition between a large number of species. We repeated this community assembly experiment from five separate pools of 600 species each. **A,B)** The metabolic networks in the final communities obtained from two separate species pools are identical. From all possible reactions (gray arrows), the same subset of reactions is active (i.e., carry a positive flux) in the final communities (colored arrows). On each reaction arrow, the size of the circle shows the flux carried by this reaction and the pie-chart shows how this flux is shared between species. The manner in which species contribute to the flux differs between pools. **C)** The fluxes through the community metabolic network are similar across communities from different species pools. The theoretical prediction (Eq. (7)) shown on the right matches simulations. The reactions in the bar plot are colored according to the arrow colors in panels A and B. **D)** The relative abundance of species in the final communities. The colors of species in pools 1 and 2 match their colors in the pie-charts in panels A and B respectively. The relative abundance is not proportional to the flux catalyzed by the species because the energy-yield per unit flux differs between reactions. For example, the green species from pool 1 (panel A) and the pink species from pool 2 (panel B) catalyze a large flux but have a low relative abundance due to the low energy yield of the reaction *R*_0_ *→ R*_1_ (panel C).

#### Functional convergence

We compared the metabolic functioning of the communities obtained from different species pools. The function performed by the community was defined as the flux through the active reactions in the final community. Individual reaction fluxes, visualized as radii of circles in Fig. 2A,B and color bars in Fig. 2C, were highly similar across pools. Thus, communities converged in the metabolic function they performed despite having been assembled from different species pools.

#### Taxonomic divergence

Bolstered by the convergence in metabolic network structure and function across pools, we measured abundances and functional roles of individual species in communities assembled from different species pools. The functional role of a species was quantified as the reaction flux catalyzed by it. We found that species contributed to the reaction fluxes idiosyncratically across pools (pie charts in Fig. 2A,B). Several reactions were performed by multiple species while others were performed by only one, and this breakup of reaction fluxes between the species varied between pools. The relative abundance of a species (Fig. 2D) is determined by the sum of all metabolic fluxes catalyzed by its enzymes multiplired by the energies assimilated per unit flux, which differed between reactions. Unlike community fluxes through individual metabolic reactions, relative abundances and functional roles of individual species did not converge across different pools.

Taken together, the three observations above mirror the functional convergence despite taxonomic divergence observed in many natural communities^30–32^. We investigated the model analytically to understand why these slow-growing, energy-limited communities share these emergent properties. We uncovered fundamental thermodynamic principles governing functional convergence in these energy-limited communities, which we explain in the next section.

### Principle of maximum free energy dissipation explains selected community metabolic network

We first sought to understand why the same metabolic reactions are realized in the final community across species pools. We were able to make analytical progress by focusing our attention on slow-growing communities growing on low-energy reactions. At steady-state, the biomass growth in communities replaces loss by dilution, *δ*. In slow-growing communities, the growth at steady-state is much smaller than the maximum growth rate of the species *g*_max_. Motivated by conditions in ocean sediments and bioreactors, the substrate concentration in the resource supply in simulations was larger than the typical *K_S_*^3, 30, 48, 51^. The high substrate levels in the supply mean that growth was primarily slowed by the accumulation of products. Since growth is slowed down by product accumulation, we can relate the concentrations of substrates and products of an active reaction in the final steady-state community. As an example, consider reaction *R*_0_ *→ R*_1_ in a community with four resources (Fig. 3A). If the reaction is active in the final steady-state community, then we have:

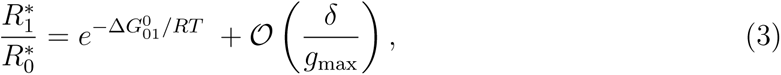

**FIG. 3.**
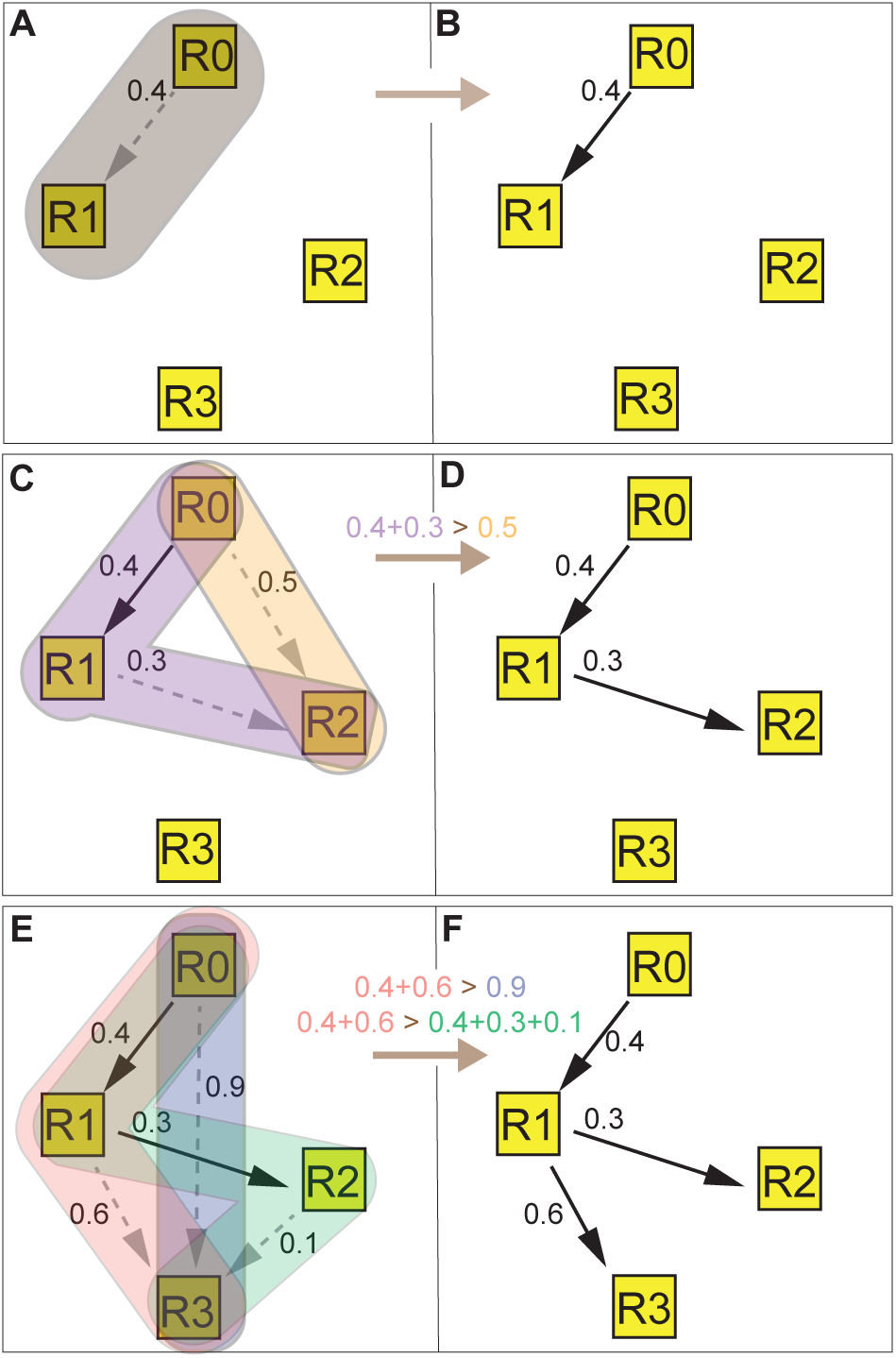
The maximum dissipation principle determines the community metabolic net-work. In a system with four resources, we assemble the community metabolic network by con-sidering the production of each resource sequentially to understand the governing thermodynamic principles. **A)** The sole path from the supplied resource, *R*_0_, to the next-highest energy resource *R*_1_. The path dissipates 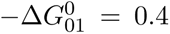 units of energy. **B)** The steady-state community with reaction *R*_0_ *→ R*_1_ maintains resource concentrations such that 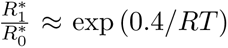. **C)** The two possible paths from *R*_0_ to the next resource, *R*_2_, that could be added to the community. The paths *R*_0_ *→ R*_2_ and *R*_0_ *→ R*_1_ *→ R*_2_ dissipate 0.5 and 0.7 units of energy. **D)** The path that dissipates more free energy, *R*_0_ *→ R*_1_ *→ R*_2_, is selected in the final community because species utilizing this path will drive concentration of *R*_2_ higher (such that 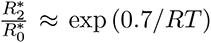) than species utilizing *R*_0_ *→ R*_2_ **E)** The three possible paths from *R*_0_ to *R*_3_, that could be added to the community. We do not consider the path *R*_0_ *→ R*_2_ *→ R*_3_ because we have already found that the maximally dissipative path to *R*_2_ is *R*_0_ *→ R*_1_ *→ R*_2_ and not *R*_0_ *→ R*_2_. **F)** The path that dissipates most free energy, *R*_0_ *→ R*_1_ *→ R*_3_, is selected in the final community because it can drive the concentration gradient *R*_3_*/R*_0_ highest (such that 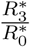 *≈* exp (10*/RT*)). Thus, the active reactions in the final community network will be determined by the paths to each resource that dissipates the most free energy.

where 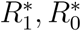 are the steady-state concentrations of the resources and 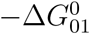 is the free energy dissipated in the reaction *R*_0_ *→ R*_1_, defined as the negative of the standard free energy difference of the reaction (Eq. (1)). This condition implies that reactions are slowed down to near thermodynamic equilibrium due to product accumulation. The first (and leading order) term depends only on the thermodynamics of the reaction. The second term, 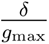, which contains all the properties specific to the species and enzyme, is only a small correction.

We now consider the production of the second resource, *R*_2_, which can be produced via two possible reaction paths (Fig. 3C). To analyze this scenario, we derive a crucial extension to Eq.(3) that applies to reaction paths, composed of multiple reactions, that are active in the final community (see SI for derivation). If a reaction path proceeding from *R_i_* to *R_j_* is active in the final community, we have:

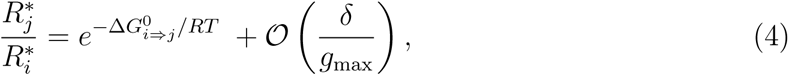

where 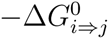 refers to the sum of the free energy dissipated in the reactions along the path from *R_i_* to *R_j_*. Again, note that the species-and enzyme-specific properties are only a small correction. This equation provides us different values for 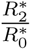 for each of the paths to *R*_2_ considered (Fig. 3C). Since both values cannot hold simultaneously, only one of the two paths can be realized in the final community. Thus, the in-degree of any node in the network is always one (c.f. Fig. 2A,B).

To understand which of the two paths is selected in the final community, we can consider the competition of two species utilizing reactions *R*_1_ *→ R*_2_ and *R*_0_ *→ R*_2_ added to the community with reaction *R*_0_ *→ R*_1_ (see Fig. 3B). The species utilizing the reaction *R*_1_ *→ R*_2_ (and hence path *R → R → R*) will drive product accumulation until 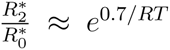, while the species utilizing the reaction *R*_0_ *→ R*_2_ will drive product accumulation until 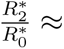 *e*^0^.^5^*^/RT^*. Since *e*^0^.^7^*^/RT^ > e*^0^.^5^*^/RT^*, the former species can invade and grow in the steady-state environment of the latter, driving the product accumulation to levels where the latter cannot grow. Thus, the species utilizing the path dissipating more free energy is able to invade and displace its competitor, and create conditions that cannot be invaded by the other. Hence, the path to *R*_2_ that dissipates more free energy is realized in the final community (Fig. 3D).

The same argument can be extend to resources with multiple paths leading to them, such as *R*_3_. We consider three paths to *R*_3_ (Fig. 3E). Note that we exclude the path *R*_0_ *→ R*_2_ *→ R*_3_ since we previously found the maximally dissipative path to *R*_2_; this is an application of Prim’s algorithm to find the maximal spanning tree in a network^52^. Comparing the free energy dissipated among the three paths, we find the maximally dissipative path, *R*_0_ *→ R*_1_ *→ R*_3_. A species utilizing this path is able to raise the concentration of *R*_3_ high enough so as to make the other reactions infeasible, driving competitors relying on the paths extinct. Thus, the principle of maximum dissipation governs the selection of the community metabolic network in slow-growing, energy-limited communities.

### Functional convergence from thermodynamic principles

The maximum dissipation principle explains why the same reactions are realized in the final communities across pools. But it does not explain the convergence in reaction fluxes across pools.

To understand how this convergence in fluxes arises, we build on results in Eq. (4) to derive the concentration at steady state of *R*_0_,

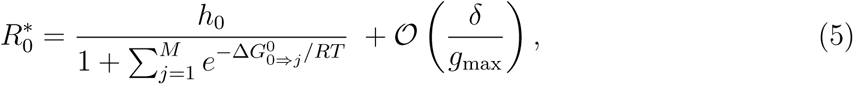

where *h*_0_ is the steady state concentration of *R*_0_ in absence of any consumption and 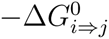 is the free energy dissipated in the reaction path to *R_j_* realized in the community (see SI). Using Eq. (4), we can obtain the the steady-state concentrations of all resources:

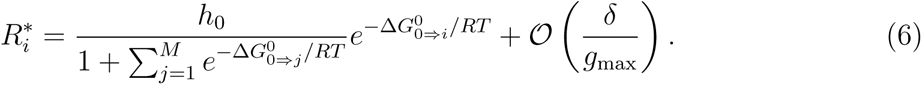

These equations imply that the resource concentrations at steady-state are determined to the leading order by thermodynamics alone. The surviving species grow to abundances that allow them to maintain the environmental resource concentrations at these thermodynamically determined values. Note that these equations resemble the Boltzmann-Gibbs distribution in statistical physics, now realized in an ecological context.

The fluxes in the steady-state community can be calculated by conserving resource flux along the chemical reaction network. The outflow from a terminal resource without any outgoing reactions, *R_T_*, is solely determined by dilution and equal to 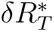. At steady state, this outflow is balanced by the reaction flux into the terminal resource. This can be used to calculate the flow from resources producing a terminal resource. Iterating this procedure, we can calculate the total flux into any resource *R_i_*, *I_i_*, as

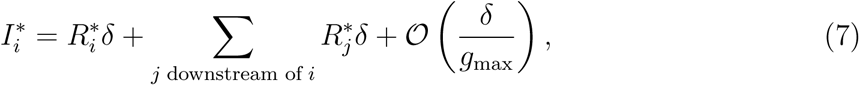

where the sum is over all resources, *R_j_*, that appear downstream of *R_i_* in the reaction network. This theoretical prediction closely matches simulations (Fig. 2C), with a small deviation due to the finite dilution rate in simulations. Importantly, the steady-state resource concentrations and reaction fluxes are independent of enzyme budgets, enzyme allocation, and details of the chemical kinetics—none of the associated parameters (*E, k_cat_, K_M_, K_S_*) appear in the equations to leading order (Eq.6). Hence there is strong selection for choosing the maximally dissipative network and converging to the thermodynamically constrained fluxes; selection on the species-specific properties is weaker. To summarize, we observe functional convergence, driven by the shared reaction thermodynamics across experiments, despite taxonomic divergence, in terms of how species conspire to maintain the community function.

Note that a community utilizing the maximally dissipative reactions can still experience species turnover. For example, an invading species with larger proteome allocation to the set of maximally dissipative reactions can displace a resident species. However, this invasion will have only a very small effect (of order 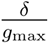) on the community metabolic function and steady-state resource environment. Hence, communities can continuously experience species turnover while maintaining their metabolic functions, as seen in bioreactors^30, 33^.

### Functional convergence decreases with community growth rate

An important assumption in our theoretical derivation of the maximal dissipation principle and thermodynamic explanation of functional convergence was that communities were slow-growing. But what happens when growth rate increases? Our theoretical analysis predicts that differences in species’ enzyme budgets and allocation, and in enzymatic parameters, will start to matter as growth rate increase, and will cause greater differences in the community metabolic network structure and function of communities from different pools. To study community metabolic network structure and function as the steady-state growth rate increases, we simulated community assembly from 10 separate pools of 600 species at different dilution rates *δ*. Since the growth rate of a community at steady-state is equal to the dilution rate, these simulations provided us with 10 different final communities at each steady-state growth rate.

Fig. 4A,B,C depicts three final communities obtained, at three different dilution rates from the same species pool. The maximum dissipation principle predicts all five reactions correctly in the slow-growing community, which grows at 1% of the maximum possible growth rate (dashed lines indicate reactions predicted correctly). At an intermediate growth rate, the maximum dissipation principle predicted three out of five reactions correctly. Finally, for the fast-growing community, which grew at 30% of the maximum possible growth rate, only two out of five reactions were predicted correctly. The error in the prediction by the maximum dissipation principle, defined as the fraction of reactions that disagree with the predictions, averaged across the 10 species pools, increases with the community growth rate (Fig. 4D). Communities deviate from our predictions at higher dilution rates due to a combination of factors. First, due to the low energy yields of reactions on the maximally dissipative path, species utilizing these reactions need to catalyze a larger flux to achieve high growth rates than species utilizing reactions with higher energy yields on competing paths. Second, due to the variation in enzyme budgets and enzyme allocation strategies, a species utilizing reactions on a maximally dissipative path can get out-competed by a species with a larger enzyme budget allocated to reactions on a competing path. Third, due to the variation in enzymatic parameters, a reaction on a competing path might be preferred because it has better enzymes, (e.g. with higher *k*_cat_).

**FIG. 4.**
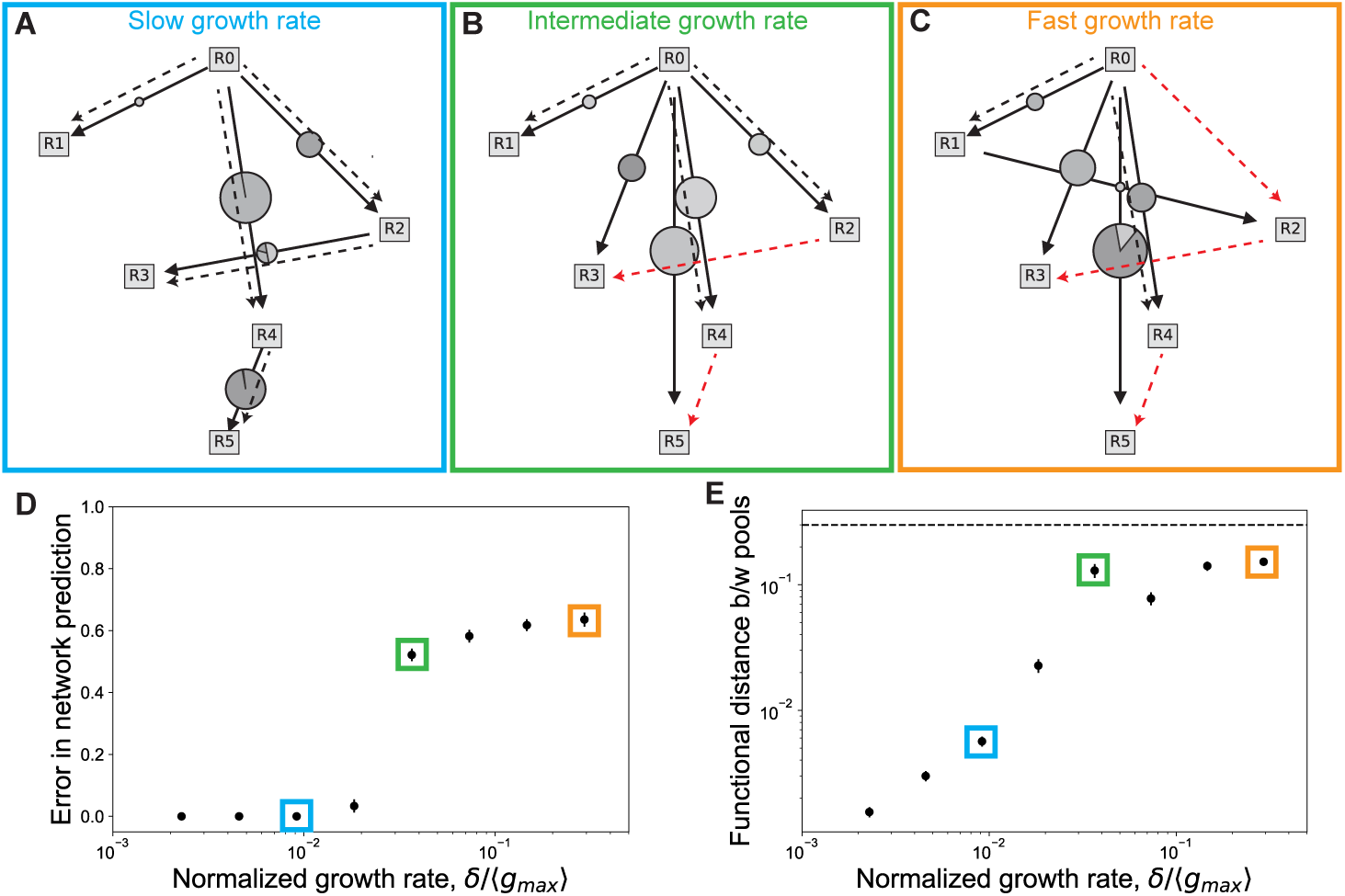
Functional convergence and the strength of thermodynamic constraints decreases in fast-growing communities. **A),B),C)** The realized community metabolic networks at slow growth rate, intermediate growth rate, and fast growth rate respectively. All three communities were obtained from the same species pool, and the steady-state growth rate was varied by changing the dilution rate, *δ*. Solid lines indicated realized reactions; dashed lines represent predictions based on the maximum dissipation principle; red lines indicate reactions that disagree with the predictions. In the slow-growing community, the active reactions match the five reactions predicted from the maximum dissipation principle; in the middle community, three of the active reactions match the prediction; in the fast-growing community, only two of the active reactions match the prediction. **D)** The fraction of reactions that disagree with the prediction of the maximum dissipation principle, averaged across 10 different species pools in the same environment. The error in the predicted network increases with the community growth rate. **E)** The functional distance between two communities was quantified by the Jensen-Shannon distance between the two community flux vectors. The average functional distance increases with the growth rate. Thus, the observed functional convergence decreases as the communities grow faster. The dashed line indicates the average distance between two random flux vectors. A random flux vector had the same length (15), and had each element picked from a uniform distribution between 0 and 1. The colored boxes in panels D and E correspond to the growth rates shown in the panels in A,B,C. The maximum growth rate, *g_max_*, was used to normalize the steady-state growth rate. *g_max_* was defined as the growth rate obtained on an irreversible reaction between consecutive resources by a species investing its entire budget in the reaction without any energy dissipation (see Methods).

As the idiosyncratic properties of species pools and enzymes start to play a bigger role in determining the community metabolic network, functional convergence weakens. The average distance between community fluxes (i.e., the degree of functional divergence) increases in fast-growing communities (Fig. 4E). The dashed line shows the expected distance between two random flux vectors; this distance is higher than the most functionally distant communities because the random flux vectors are not subject to ecological and flux constraints. Together, these observations show that the maximum dissipation principle and functional convergence will be stronger in low-energy communities that grow slowly, yet some degree of convergence is expected to persist even for faster growth rates.

### Validating predictions using experimental data from anaerobic digesters

We wanted to validate some of our predictions using real-world experimental data. Anaerobic digesters present a good test system because the lack of strong electron acceptors forces microbes to use low-free energy reactions to fuel growth. The study by Peces et al.^53^ examined the metabolic function of four anaerobic digesters, originally inoculated with four different microbial communities. Each digester was run at four dilution rates (hydraulic retention times of 15, 8, 4, and 2 days), and metabolic concentrations were measured at steady-state for each dilution rate. The reported data allowed us to compare model predictions with experimental data on how functional convergence varied with dilution rate.

In line with our predictions, we observed convergent metabolic concentrations in the four digesters at low dilution rates, and metabolic convergence weakened with increasing dilution rate (Fig. 5). Specifically, the variability in measured metabolic concentrations across the four digesters increased at higher dilution rates, whether quantified for each metabolite individually (Fig. 5 A) or together (Fig. 5 B). Simulations of four species pools at different dilution rates reproduced this decrease in metabolic convergence with increasing dilution rate (Fig. 5 C,D). Furthermore, a comparison of other published results suggests that this loss of functional convergence holds more generally, across studies. The functional stability of communities in anaerobic bioreactors was measured in three studies, each at a different dilution time (15, 10 and 1 days)^30, 33, 54^. While the two studies at the lower dilution rates observed functional convergence, the study at the highest dilution rate did not, in agreement with our predictions. Thus, our results can explain not only functional convergence in microbial communities, but also the conditions under which it may disappear.

**FIG. 5.**
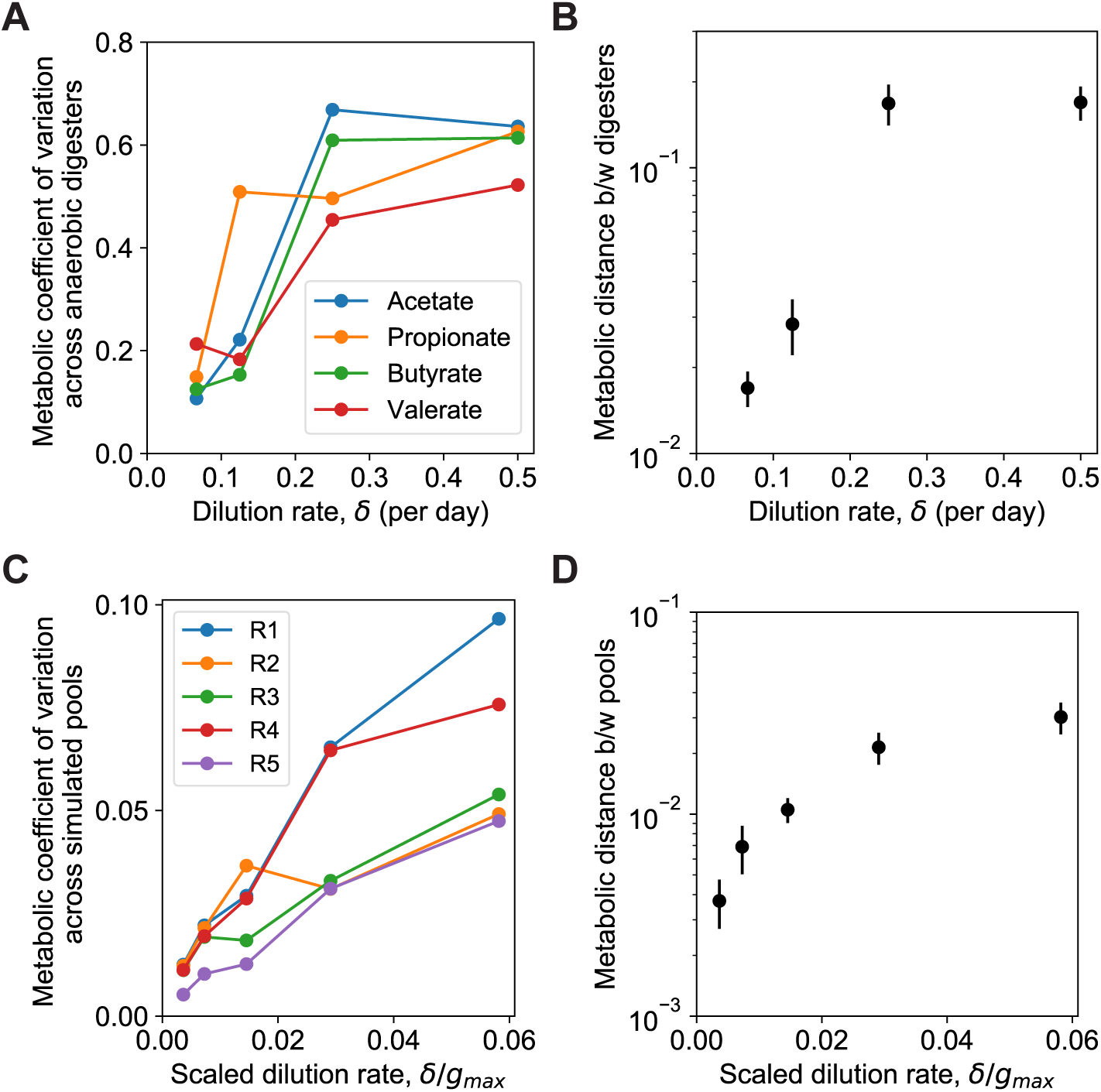
Metabolic convergence in experimental anaerobic digesters weakens with dilution rate. We measured the metabolic convergence across four anaerobic digesters using data from Ref.^53^. In line with model expectations, the variability in resource concentrations across the four digesters increased with dilution rate. **A)** The coefficient of variation (ratio of standard deviation to mean) of organic acid concentrations across four anaerobic digesters increases with the dilution rate of the digester. **B)** The metabolic distance between digesters, quantified by the average Jensen-Shannon distance between resource concentrations in the digesters. **C,D)** Simulations of our model with four species pools at various dilution rates showed a similar increase in the metabolic coefficient of variation and metabolic distance with dilution rate. The input flux of resource in simulations was held constant, as in the experiment.

### Functional convergence weakens but persists in higher-energy environments

All of our results apply to microbial communities growing in low-energy environments, where catabolic reactions are reversible. But what happens when species grow in environments with at least some of the higher-energy reactions being effectively irreversible? Our theoretical analysis predicts that as reactions become less reversible, differences in species’ enzyme budgets, allocation, and kinetic parameters start to matter, increasing the differences in the metabolic network structure and function of communities from different pools. To study changes in community metabolic network structure and function as reaction energies increase, we simulated its assembly from four separate pools of 600 species in environments characterized by different reaction energies. Specifically, to obtain environments in which reactions progressively become less reversible, we scaled all the energies in the system: the energy of each resource, the energy assimilated in each reaction, and the energy dissipated in each reaction by a common factor (see Methods for further details). We record the results as a function of the standard reaction energies in the system since it is the most easily available quantity for real communities.

We explored the community metabolic network structure as the reaction energies in the environment were varied. Fig. 6A depicts communities obtained from three different species pools in a low-energy environment where the energy gap in reaction *R*_0_ *→ R*_5_ was 5*RT* (12.4 kJ*/*mol). In this scenario, all communities converged to the network predicted by maximum dissipation (indicated by dashed lines). Fig. 6B depicts communities obtained from three different species pools in a higher-energy environment where the standard energy gap in reaction *R*_0_ *→ R*_5_ was 25*RT* (60 kJ*/*mol). At this higher energy, the theoretical predictions start to disagree with the realized community network (disagreements are highlighted in red). Despite these disagreements, 65% of the reactions are predicted correctly even at this higher energy.

**FIG. 6.**
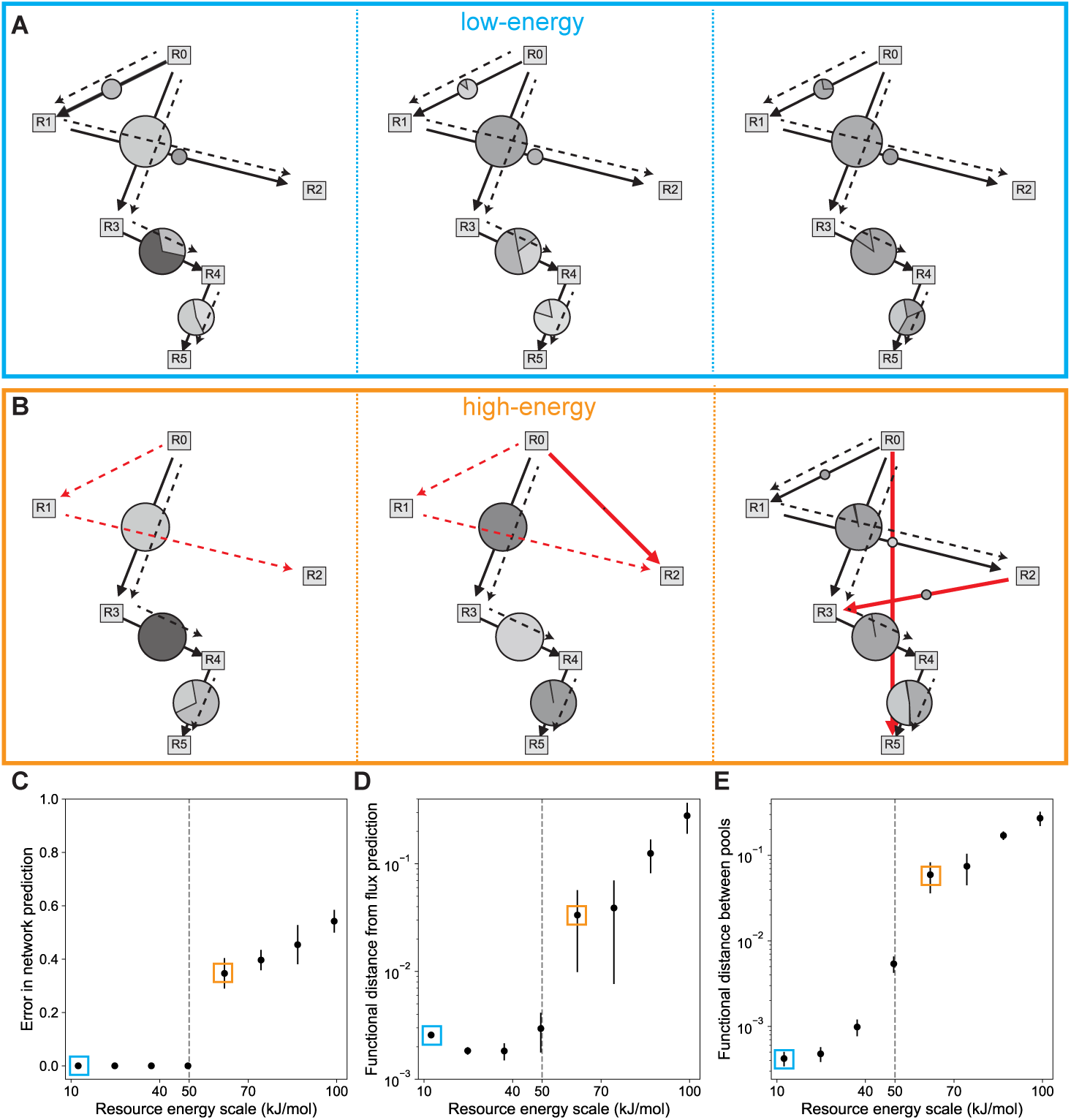
Functional convergence decreases with increasing reaction energies. Examples of realized community metabolic networks from three different species pools in an environment with low-energy reactions **(A)** and an environment with high-energy reactions **(B)**. All reaction energies are increased proportionally in these two environments. Solid lines indicate realized reactions; dashed lines represent predictions based on the maximum dissipation principle; red lines indicate reactions that disagree with our predictions. **C)** The fraction of reactions that disagree with the prediction of the maximum dissipation principle starts to increase once reaction energies cross a threshold around 50 kJ/mol (vertical line). **D,E)** The functional distance between observed and predicted fluxes as well as between observed fluxes in two different communities increases as resource energies increase. The x-axis indicates the energy of reaction *R*_0_ *→ R*_5_. Data points enclosed in blue and orange boxes correspond to parameters used to simulate low-energy (panel A) and high-energy (panel B) environments respectively. Functional distance was quantified by the Jensen-Shannon distance between community flux vectors (see Methods).

We quantified the degree of deviation from theoretical predictions using two separate measures. First, we quantified the network prediction error as the fraction of observed reactions that disagreed with theoretical predictions (Fig. 6C). Then, we quantified functional differences between realized and predicted fluxes using the Jensen-Shannon distance (Fig. 6D). Across both measures, we observed that theoretical predictions were highly accurate up to energies around 20*RT* (50 kJ*/*mol). We also quantified the degree of functional convergence in the observed communities, independent of the theoretical predictions, using the average Jensen-Shannon distance between the fluxes observed in the communities assembled from different species pools (Fig. 6E). We observe that functional convergence is progressively lost as we cross the same energy scale around 20*RT* (50 kJ*/*mol). Incidentally, this energy scale matches the energy scale up to which we expect product inhibition to be relevant in real environments, as estimated from experimental parameters (see Discussion and SI Sec. S9).

We also tested the effect of increasing the energy dissipated in specific reactions one at a time. In this study, we simulated separate experiments setting the energy dissipated in a specific reaction to a larger value. This made the chosen reaction significantly more irreversible (see SI Sec. S8). Although making any single reaction irreversible can generally cause deviations from our predictions and decrease functional convergence of the community between species pools, we found the accuracy of the predicted network to still be greater than expected from a null model irrespective of which reaction we modified. Further, we found that making reactions closer to the top of the network irreversible was less disruptive to our predictions (see SI Fig. S7). This may resemble the situation in some anaerobic digesters fed with high-energy substrates, where reactions near the top of the network have larger free energy gaps than reactions near the bottom^8^.

## DISCUSSION

Microbial community models have successfully exposed the organizational principles and universal, emergent behavior of microbial communities structured by nutrient limitation^18–24^. However, in many natural and industrial environments, microbial communities grow under energy-limited conditions and are forced to use low free-energy reactions, such as anaerobic respiration using sulfate or carbon dioxide and certain fermentation reactions, to drive growth^3–14^. Here, by explicitly modeling the thermodynamic inhibition of microbial catabolism and energy-limitation of growth, we have developed a minimal model of microbial communities in low-energy environments. We used this model to investigate the emergent properties of slow-growing microbial communities assembled in low-energy environments. Notably, our simulations displayed functional convergence despite taxonomic divergence observed in many real communities^30–32^. We demonstrated that functional convergence in our model originates from the fundamental thermodynamic constraints on microbial growth despite differences in metabolic capabilities of surviving species. This should be contrasted with previous models, where the emergence of functional convergence in nutrient-limited communities is explained by shared metabolic capabilities of all species^40^. Further, we demonstrated that community metabolic network structure in slow-growing communities assembled by ecological competition over long periods of time is determined by the thermodynamic principle of maximum free energy dissipation. Importantly, network selection by maximum dissipation manifests through ecological competition and is not expected in isolated microbes. Analyzing data from anaerobic digesters, we found that functional convergence is lost at higher dilution rates as predicted by our model. Thus, our results can not only explain functional convergence in microbial communities, but also the conditions under which it may disappear.

For simplicity, we demonstrated our main results using a minimal model of energy-limited communities. In the supplementary text, we show that our results hold more generally. In particular, we demonstrate that our results apply in the following scenarios: First, if reaction stoichiometry varies beyond 1: 1 in the model (see SI Sec.6). Second, if maintenance energy requirements of the cell are incorporated in the model; the maintenance energy flux requirement resembles biomass dilution, and so our results apply as long as this requirement is small (see SI Sec.5). And third, a model without explicit dilution of resources, where cells uptake resources for anabolic processes in addition to catabolism (see SI Sec.7).

Microbes in low-energy environments face a trade-off between harvesting energy efficiently, to increase biomass yield per unit reaction flux, and dissipating more free energy, to increase reaction flux. A microbe cannot grow at either extreme—assimilating all the energy curbs growth because the reaction flux will be zero, while dissipating all the energy curbs growth because energy assimilated will be zero. Thus, microbes have to find a balance between these opposing forces. Our results suggest that, at small dilution rates, the effect of ecological competition will push microbes to evolve towards dissipating more energy. However, in practice this evolutionary drive to decrease energy assimilation will not continue forever. It will probably be limited by biochemical constraints^41, 55, 56^, such as the smallest energy unit that can be assimilated by a species (e.g. the energy generated by a single electron, or in other cases that of one molecule of ATP), or when reactions become limited by substrate depletion rather than thermodynamic inhibition. In our simulations, to focus on the ecological properties of energy-limited communities, a species at the lower-limit of energy assimilation could survive at the smallest dilution rates considered. An exploration of the evolutionary consequences of the selective pressures on energy assimilation is reserved for future work.

We have shown that in sufficiently low-energy environments, the thermodynamics of microbial growth gives rise to functional convergence. But when can an environment be considered sufficiently low-energy? To answer this question, we compared microbial growth rates in communities with and without thermodynamic inhibition. These rates are significantly different provided that 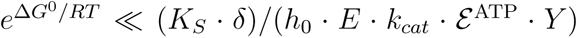 (SI Sec. S9). By obtaining estimates of the various parameters on the right hand-side of this equation from the literature (see SI Sec. S9), we were able to estimate the free energy dissipation required to make a reaction effectively irreversible. Specifically, we found that free energy dissipation *−*Δ*G*^0^ *≫* 12*RT*, which is approximately 30kJ*/* mol, is required to significantly weaken the effects of thermodynamic inhibition. Due to biochemical constraints, cells typically assimilate at least about 20kJ*/* mol from catabolic reactions^41, 55–57^. Hence for a reaction to be considered effectively irreversible, its overall energy combining dissipation and assimilation needs to be greater than about 50kJ*/* mol. Thus, an environment where the typical free energy of catabolic reactions satisfies *−*Δ*G <* 50kJ*/* mol would be affected by thermodynamic inhibition and could approximately described by our model. A fair number of natural and industrial environments fall into this category^10, 12, 13, 57, 58^.

Our model provides one possible solution to the long-standing question^30^ of why some microbial communities are characterized by functional stability in spite of taxonomic divergence. This solution can be contextualized within other recently proposed explanations. One possible explanation for functional universality in microbial communities is the identical stoichiometry of conversion between metabolic substrates and byproducts used by several consumer-resource models^39, 40^. This mechanism may explain the apparent functional stability in models where all species use the same stoichiometric matrix to convert substrates to products^40^. However, it cannot account for functional universality in microbial ecosystems where consumption of a substrate may yield different products depending on the species metabolism^8, 59^, as in our model. Indeed, different microbes in our model contain different subsets of enzymes and therefore differ from each other in how they transform nutrients in the system to byproducts. Despite this wide variation in the metabolic capabilities of the species, the system still exhibits functional stability due to universal thermodynamic principles. Other possible mechanisms for functional stability have also been proposed^60–62;^ these works describe reaction-centric mathematical models applicable to poorly mixed microbial communities, such as where metabolic rates are limited by physical transport of substrates across space. However, these mechanisms do not apply to well-mixed ecosystems, such as continuous stirred anaerobic digesters^53^.

In our model, we relate microbial growth and thermodynamics by assuming that biomass yield is proportional to energy assimilated from catabolism, which, in turn, is described by reversible Michaelis-Menten kinetics following prior experimental studies^6, 7, 44–47^. Alternative approaches relating microbial growth and thermodynamics exist^63, 64^ and have been classified in Ref^63^. Based on this proposed classification, our model and others^29, 44, 65^ fall into the first class of models, where microbial biomass yield per catabolic reaction is fixed (since yield is determined by energy assimilated in catabolism). An alternative category of models building on empirical work^66, 67^ allow biomass yield per catabolic reaction to vary dynamically based on the available energy in the environment^68, 69^. Although a complete survey of all possible models is beyond the scope of this study, we tested how this alternative assumption of a dynamic biomass yield affects our results by incorporating it into our model (see SI Sec. S10). We found our main results of functional convergence and maximum dissipation were qualitatively unaffected in a model incorporating the dynamic yield approach (see SI Fig. S8).

Optimization principles grounded in thermodynamics have been used in microbial metabolism to explain behavior at the cellular scale^70–72^. At the community level, optimization of thermodynamic quantities has been conjectured to be a mechanism by which communities self-organize, motivated mostly by verbal arguments and analogies with physical systems^73, 74^. However, the question of which quantity should be maximized, and under what conditions, remains actively disputed in the literature^34, 75^. By incorporating ecological and thermodynamic principles, our model connects these two disparate approaches toward understanding microbial communities. We derive the thermodynamic optimization principle of maximum dissipation, which structures slow-growing, energy-limited communities, providing a concrete example of thermodynamic optimization in ecology.

Our results make predictions for the metabolic structure and resource environment created by the communities in low-energy environments, based on the thermodynamics of microbial metabolism. This requires knowledge of the free energy differences and metabolomic data of the resource environment. Although calculating free energies of many metabolites present in real systems is challenging, recent advances in computational methods make the outlook promising^76–78^. Combining these methods with metabolomic data and knowledge of microbial metabolism to predict metabolite composition in real environments shaped by microbial communities is an exciting prospect for future research.

## METHODS

### Species and resource dynamics

Species dynamics are determined by the amount of reaction flux they can utilize, the energy they assimilate from this to create biomass, and biomass loss due to dilution. The dynamics of species *α* with abundance *N_α_* is described by

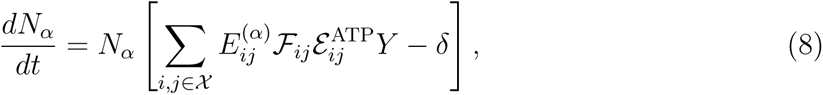

where *F_ij_* is the flux of the reaction *R_i_ → R_j_* catalyzed by a single enzyme, 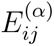 is the number of enzymes allocated to this reaction by the species *α*, *Y* is the biomass yield, *X* is the set of all allowed reactions, and *δ* is the dilution rate. The maintenance energy requirements of a species can be accounted for by replacing *δ* with *δ* + *m* in Eq. 8, where *m* is the energy flux required for cell maintenance. Our results apply in this scenario as well (see SI Secs. S5,S7). Resource dynamics are determined by their consumption and production via the reactions catalyzed by the species, external supply, and dilution. The dynamics of resource *i* with concentration *R_i_* are given by

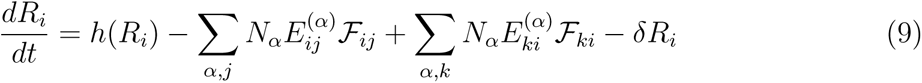

where *h*(*R_i_*) is the resource supply. Only the top resource *R*_0_ is supplied, so, *h*(*R*_0_) = *δ · h*_0_ and zero otherwise. *h*_0_ is the concentration of resource in the supply. The second and third terms describe consumption and production fluxes of the resource by the species, with *j* indexing the resources that could be produced from *i* and *k* indexing the resources that could produce *i*.

### Simulation procedure

All species in a pool were introduced at the same initial density (=1) in an environment with resource concentration of the supplied resource *R*_0_ = *h*_0_ and all other resources at a very small concentration 10*^−^*^10^. The species, reaction, and resource dynamics were simulated till they reached a steady-state. At regular intervals, species that fell below an abundance threshold had their concentration set to zero. The minimum allowed resource concentration was 10*^−^*^10^ to prevent division by zero in Eq. (2). A steady-state was reached if the maximum value of the vector 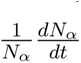, measuring the logarithmic growth rates, fell below 10*^−^*^5^. Simulations starting from different initial species concentrations also converged to the same final steady-state. Simulations were performed in python and numerical integration used the SciPy package^79^.

### Simulation parameters

The enzyme budget of a species was chosen from a lognormal distribution with lognormal parameters *µ* = 1*, σ* = 0.1. A species utilized a random subset of the 15 possible reactions. So species in the pool were not biased towards extreme generalists or specialists. The enzyme budget was allocated to the selected reactions uniformly, by using a Dirichlet distribution with the Dirichlet parameter, *α* = 1, for the selected reactions. There were 600 species in each of the 5 pools in Fig. 2 and 10 pools in Fig. 4; species were not shared across pools. In Figs. 2,4 the energies of resources *R*_0_, *R*_1_, *R*_2_, *R*_3_, *R*_4_, and *R*_5_ are respectively 5*RT*, 4*RT*, 3*RT*, 2*RT*, 1*RT,* and 0*RT*. For each reaction, a random fraction between 15% and 85% of the standard energy gap between product and substrate 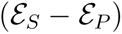 was assimilated, and the rest was dissipated. This assimilated energy, along with resource energies, was fixed across the experiments with different species pools. The concentration of *R*_0_ in the supply, *h*_0_ = 80. The chemical kinetic parameters *K_P_*,*K_S_*, and *k_cat_* were chosen from lognormal distributions with lognormal parameters *µ* = 0*, σ* = 0.01, and so, their mean was close to 1. The energy to biomass yield factor, *Y* = 1. The dilution rate, *δ* = 0.01 in Fig. 2; this rate was small enough that a typical species would survive if it was alone.

In Fig. 5, the input flux of *R*_0_ was held fixed across dilution rates to mimic the experiment (*R*_0_*δ* = 10). We simulated four different species pools, starting with 600 species in each pool. The energies of resources *R*_0_*, R*_1_*, R*_2_*, R*_3_*, R*_4_, and *R*_5_ were 5*RT,* 4.2*RT,* 3.7*RT,* 2.4*RT,* 1.4*RT,* and 0*RT*. Motivated by real-world scenarios, the energy gaps were not uniformly spaced to demonstrate the robustness of our results to differences in spacing. For the same reason, we increased the degree of variation in chemical kinetic parameters *K_P_*,*K_S_*, and *k_cat_*, which were chosen from lognormal distributions with lognormal parameters *µ* = 0*, σ* = 0.15. The concentration of *R*_0_ in the supply *h*_0_ = 80 and energy to biomass yield factor, *Y* = 1.

In Fig. 6, the energies of resources *R*_0_*, R*_1_*, R*_2_*, R*_3_*, R*_4_, and *R*_5_ were selected to be proportional to 5*RT,* 4*RT,* 3*RT,* 2*RT,* 1*RT,* and 0*RT*. The resources energies and the energies assimilated and dissipated in individual reactions were scaled by the same factor to generate different environments. The energy of *R*_0_, which is plotted on the x-axis; is also the energy gap in the reaction *R*_0_ *→ R*_5_. The concentration of *R*_0_ in the supply line to the chemostat is *h*_0_ = 80. For each reaction, a random fraction between 40% and 80% of the standard energy gap between product and substrate was assumed to be assimilated into reusalble energy carries such as ATP (based on experimental values^65,80^and the rest was dissipated. The chemical kinetic parameters *K_P_*,*K_S_*, and *k_cat_* were chosen from lognormal distributions with lognormal parameters *µ* = 0*, σ* = 0.01. The energy to biomass yield factor, *Y* = 1, and dilution rate, *δ* = 0.01.

The Jensen-Shannon distance (JSD) is used to measure functional distance between two flux vectors in Fig. 4 and concentration vectors in Fig. 5. The JSD is the square root of the symmetrized Kullback-Leibler divergence, and is defined as

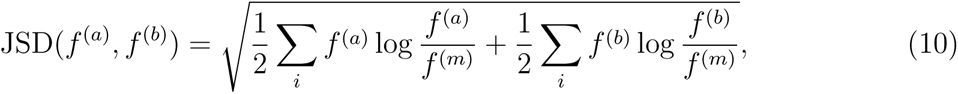

where *f*^(^*^a^*^)^*, f* ^(^*^b^*^)^ are the two normalized flux vectors and *f* ^(^*^m^*^)^ is an average flux vector defined as (*f* ^(^*^a^*^)^ +*f* ^(^*^b^*^)^)*/*2 (see Refs.^79, 81^). We report JSD values averaged over all pairs of communities in Figs. 4, 5. Flux arrows in the metabolic network are plotted only if the relative flux in the reaction is *>* 10*^−^*^4^.

The error in the network predicted by maximum dissipated was calculated as 1*−*accuracy. The accuracy was defined as the fraction of reactions correctly predicted:

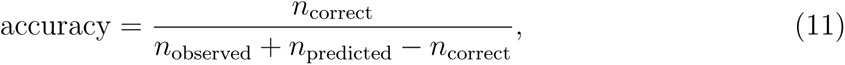

where *n*_observed_ is the number of reactions in the observed network, *n*_predicted_ is the number of reactions in the predicted network, and *n*_correct_ is the number of reactions correctly predicted. The numerator can be understood as the number of elements in the intersection of the predicted and observed sets of reactions, while the denominator is the number of elements in the union of the two sets. If the sets are identical, accuracy is one and error is zero.

*g_max_* in Figs. 4, 5 was defined as the growth rate obtained from an irreversible reaction where a species invested its entire enzyme budget into one reaction. Since this varies between reactions because reactions have different energy yields, we chose the energy gap between consecutive resources. It was calculated as the product of mean *k_cat_*, mean enzyme budget, average energy gap between consecutive resources (= *RT*), and biomass yield factor *Y*.

### Experimental data

Fig. 5 analyzes experimental data^53^, which were obtained from four anaerobic digesters (continuous stirred tank reactors) run under identical conditions. The digesters were inoculated from four different sources: slaughterhouse waster, digested sewage sludge, digested pig manure, and anaerobic granules treating brewery waste water; hence the digesters differed in their taxonomic composition. The organic loading rate was held fixed while decreasing the Hydraulic Retention Time (dilution rate) in sequential steps. The dilution rate was changed only after at least three cycles of steady-state data were obtained. We used the measured mean steady-state concentrations for each digester of the four organic acids: acetate, propionate, butyrate, and valerate. Note that we ignore the methane measurements because it is a gas and was also measured by a different technique. Nevertheless, methane also showed an increasing coefficient of variation with dilution rate, in line with predictions.

## Code and data availability

Code used for simulation and analysis can be found at https://github.com/maslov-group/Thermodynamics_functional_convergence.git. Experimental data analyzed was obtained from Ref.^53^.

## Acknowledgements

The authors thank Avi Flamholz for valuable comments and feed-back on the manuscript. This research was supported in part by NSF Grant No. PHY-1748958 and the Gordon and Betty Moore Foundation Grant No. 2919.02.

## Author Contributions

All authors helped devise the study; A.B.G. performed the research and analyzed the data; S.M. supervised the research; A.B.G. wrote the manuscript with help from S.M. and T. W.

