## Supplementary material for "Functional convergence in slow-growing microbial communities arises from thermodynamic constraints": SI

April 6, 2023

#### Contents

**S1 Model** 1

**S2 Solution for the steady-state community** 3

**S3 Selection of metabolic network: principle of maximum dissipation** 7

**S4 Convergence of metabolic fluxes** 9

**S5 Incorporating maintenance energy requirements** 9

**S6 Generalising to arbitrary reaction stoichiometry** 9

**S7 A model without resource dilution** 13

**S8 Modifying the energy dissipation of individual reactions in the network** 14

**S9 How much free energy dissipation makes a reaction irreversible?** 17

**S10 An alternate prescription for relating thermodynamics and yield** 18

#### S1 Model

We consider a community model where species grow by harvesting energy from chemical reactions converting metabolic substrates into products. There are $M+1$ resources or metabolites that we label in the order of decreasing energy as $R_0, R_1, ..., R_M$, with energies $\mathcal{E}_0, \mathcal{E}_1, ..., \mathcal{E}_M$ such that $\mathcal{E}_0 > \mathcal{E}_1 > ...\mathcal{E}_M$. Species utilize reactions that convert a substrate of a higher energy into a product of lower energy. This mirrors the trend in catabolism of degrading more complex and energy-rich molecules, like glucose into simpler ones, like acetate.

Chemical reactions connect all pairs of resources, making the space of possible reactions a fully connected network with $(M+1)M/2$ possible links. From each unit flux of reaction $R_i \to R_j$ , species harvest $\mathcal{E}_{ij}^{\mathrm{ATP}}$ of energy, stored in ATP or other forms. The species convert harvested energy into biomass based on a yield factor $Y$ which is constant across species. For a species to utilize a reaction, it needs to have the corresponding enzyme. Each species allocates an overall enzyme budget $E^{\text{total}}$ among reactions it can catalyze. We allow overall enzyme budgets to vary between species to avoid any special degenerate behavior [1].

Reversible Michaelis-Menten kinetics describes reaction flux We illustrate the scheme to calculate flux through the chemical reactions by using an example reaction below. A enzyme catalyzed reaction from substrate $S$ to product $P$ can be understood as following the scheme:

$$E + S \rightleftharpoons EX \rightleftharpoons E + P, \tag{S1}$$

where $E$ is the enzyme corresponding to this reaction and $EX$ is some intermediate enzyme complex. The conversion of the substrate to the product could proceed over multiple steps; this will not make a difference to our results [2]. One can also think of this reaction as a coarse-grained or lumped description of the full reaction [3, 4, 5].

The cell harvests energy, $\mathcal{E}^{\text{ATP}}$, from the reaction, which is stored in the phosphorylation of ADP, reduction of NAD, or other forms. The energy harvesting decreases the Gibbs free energy drop across the reaction. The free energy difference of the reaction under standard conditions is

$$\Delta G^0 = \mathcal{E}_P^0 + \mathcal{E}^{\text{ATP}} - \mathcal{E}_S^0, \tag{S2}$$

where $\mathcal{E}_P, \mathcal{E}_S$ are the standard-state energies of the substrate and product.

The reaction flux is described by reversible Michaelis-Menten kinetics (Eq.2 in the main text):

$$J = k_{cat} E \frac{\frac{S}{K_S}}{1 + \frac{S}{K_S} + \frac{P}{K_P}} \left(1 - \frac{P}{S}\gamma\right), \tag{S3}$$

where $k_{cat}, K_S, K_P$ are enzymatic parameters describing chemical kinetics and $\gamma = e^{\Delta G^0/RT}$. The first term, $k_{cat}E$, describes the maximal velocity of the reaction, which we call $\nu$. The second term describes enzyme binding affinities and saturation level. The final term describes inhibition due to product accumulation. If $P$ is very large, the equation predicts a negative flux due to the reversal of the reaction direction. This would cause species to lose energy from using the reaction. So we assume that species down-regulate enzyme production to prevent energy loss from driving the reverse reaction.

We will also define the reaction flux per unit enzyme, $\mathcal{F}$ as

$$\mathcal{F} = J/E = k_{cat} \frac{\frac{S}{K_S}}{1 + \frac{S}{K_S} + \frac{P}{K_P}} \left(1 - \frac{P}{S}\gamma\right), \tag{S4}$$

#### The low and high substrate flux limits

The low and high flux regimes correspond to $S^* \ll K_S$ and $S^* \gg K_S$ respectively (see [6] for a detailed discussion of the limits).

In the low flux regime, the consumption flux is proportional to substrate concentration, i.e.,

$$J = k_{cat} E \frac{S}{K_S} \left(1 - \frac{P}{S}\gamma\right). \tag{S5}$$

In the high flux regime $S^*/K_S \gg 1 + P^*/K_P$, the consumption flux is substrate saturated

$$J = k_{cat} E \left(1 - \frac{P}{S}\gamma\right). \tag{S6}$$

##### Extending notation to multiple reactions

The flux equations described above were for a single reaction. To apply these equations in the model, which has many reactions, we add subscripts to denote the reaction. The enzyme allocated by an individual of species $\alpha$ for a reaction $R_i \to R_j$ will be $E_{ij}^{(\alpha)}$. The corresponding per-capita flux will be $J_{ij}^{(\alpha)}$. The chemical kinetic constants and free energy dissipated are chemical and enzymatic properties that vary only between reactions and not between species. Thus the flux per unit enzyme is $\mathcal{F}_{ij}$, the free energy dissipated is $-\Delta G_{ij}^0$, the strength of inhibition is $\gamma_{ij}$, etc. Since the notation becomes clumsy quickly, we will suppress indices for clarity if possible.

#### Species and resource dynamics

Species dynamics are determined the amount of reaction flux they can utilize, the energy the assimilate from this to create biomass, and biomass loss due to dilution. The dynamics of species $\alpha$ with abundance $N^{(\alpha)}$ is described by (see also Eq.8 in the main text)

$$\frac{dN_\alpha}{dt} = N_\alpha \left[ \sum_{i,j \in \mathcal{X}} E_{ij}^{(\alpha)} \mathcal{F}_{ij} \mathcal{E}_{ij}^{\mathrm{ATP}} Y - \delta \right], \tag{S7}$$

where $F_{ij}$ is the flux of reaction $R_i \to R_j$ catalyzed per enzyme, $E_{ij}^{(\alpha)}$ is the enzyme allocated to reaction $R_i \to R_j$ by species $\alpha$, $\mathcal{X}$ is the set of all allowed reactions, and $\delta$ is the dilution rate. The flux in a particular reaction catalyzed by a species is zero if the species does not allocate any enzyme to it.

Resource dynamics are determined by their consumption and production via the reactions catalyzed by the species, external supply, and dilution. The dynamics of resource $i$ with concentration $R_i$ are given by (see also Eq.9 in the main text)

$$\frac{dR_i}{dt} = h(R_i) - \sum_{\alpha,j} N_\alpha E_{ij}^{(\alpha)} \mathcal{F}_{ij} + \sum_{\alpha,k} N_\alpha E_{ki}^{(\alpha)} \mathcal{F}_{ki} - \delta R_i \tag{S8}$$

where $h(R_i)$ is the resource supply. The second and third terms described consumption and production fluxes of the resource by the species, with $j$ indexing the resources that could be produced from $i$ and $k$ indexing the resources that could produce $i$.

We study the scenario where the most energetic resource, $R_0$ is supplied. The concentration of $R_0$ in the supply is $h_0$. Thus the supply function is

$$h(R_i) = \begin{cases} h_0 \delta & \text{if } i = 0 \\ 0 & \text{otherwise.} \end{cases} \tag{S9}$$

### S2 Solution for the steady-state community

In this section, we will examine the equations describing a given community at steady state. At steady state, we can set the LHS of Eq.(S7) to zero for any surviving species $\alpha$. This gives us the condition

$$\sum_{i,j \in \mathcal{X}} E_{ij}^{(\alpha)} \mathcal{F}_{ij} \mathcal{E}_{ij}^{\mathrm{ATP}} Y = \delta \ \forall \alpha \in \text{survivor}. \tag{S10}$$

By counting the number of independent equations and degrees of freedom, we can see that the number of surviving species is limited by the number of reactions. Furthermore, because species regulate enzymes to ensure they do not catalyze reaction fluxes in the reverse direction and lose energy, the number of survivors is limited by the number of active reactions (reactions that carry a positive flux).

Restricting the equations to just the surviving species and active reactions only, we can express the Eq. (S10) as a matrix equation:

$$\mathbf{E}\overrightarrow{F}Y = \delta\,\overrightarrow{1}, \tag{S11}$$

where $\mathbf{E}$ is the matrix of enzyme allocation of the surviving species in the active reactions, $\overrightarrow{F}$ is the vector of fluxes in the active reactions, $\delta$ is the dilution rate, and $\overrightarrow{1}$ is a vector of ones corresponding to each survivor.

We can solve this equation by multiplying the inverse of $\mathbf{E}$ matrix on both sides. Substituting the definition of $\mathcal{F}$ (Eq. (S4)), we get

$$\left(1 - \gamma_{ij}\frac{R_j^*}{R_i^*}\right) = \frac{\delta}{k_{cat,ij}\mathcal{E}_{ij}^{\mathrm{ATP}}Y}\frac{1 + \frac{R_i^*}{K_{S_{ij}}} + \frac{R_j^*}{K_{P_{ij}}}}{\frac{R_i^*}{K_{S_{ij}}}}\left[\mathbf{E^{-1}}\overrightarrow{1}\right]_{ij}, \tag{S12}$$

for each active reaction $R_i \to R_j$ in the community, where $*$ is used to denote the steady state concentration of the corresponding resource and $\left[\mathbf{E^{-1}}\overrightarrow{1}\right]_{ij}$ refers to the element of the vector $\mathbf{E^{-1}}\overrightarrow{1}$ corresponding to reaction $R_i \to R_j$.

We can further simplify this in the high-flux limit (Eq. (S6)) where the second term in the RHS is one. Physically, this means that we are assuming that reaction fluxes are primarily slowed down by thermodynamic inhibition rather than substrate depletion. For each active reaction in the final community, we have a relation for the steady state concentrations of the substrate and product:

$$\frac{R_j^*}{R_i^*} = \frac{1}{\gamma_{ij}}\left(1 - \frac{\delta}{k_{cat,ij}\mathcal{E}_{ij}^{\mathrm{ATP}}Y}\left[\mathbf{E^{-1}}\overrightarrow{1}\right]_{ij}\right). \tag{S13}$$

Thus $\frac{R_j^*}{R_i^*}$ is set by thermodynamics to leading order in $\delta$.

To simplify the notation, we will define $\mu_{ij}$ as

$$\mu_{ij} = \frac{\delta}{k_{cat,ij}\mathcal{E}_{ij}^{\mathrm{ATP}}Y}\left[\mathbf{E^{-1}}.\overrightarrow{1}\right]_{ij} \tag{S14}$$

To gain intuition we consider the expression when species are specialists. In this case we have:

$$\mu_{ij} = \frac{\delta}{k_{cat,ij}\mathcal{E}_{ij}^{\mathrm{ATP}}YE_{ij}^{(\beta_{ij})}}, \tag{S15}$$

where $\beta_{ij}$ is the species specializing in reaction $R_i \to R_j$. We see that $\mu_{ij}$ compares the steady-state growth rate of the species in the community ($\delta$) to the maximum growth rate of the species, achieved in an environment without thermodynamic inhibition. Thus $\mu$ is a measure of the degree of thermodynamic inhibition of the species in the community. The specialist approximation can also be of practical use for obtaining rough estimates since we need not have access to the enzyme allocation matrix of a natural community.

In terms of $\mu_{ij}$, the steady-state concentration of the substrate and product in an active reaction is given by

$$\frac{R_j^*}{R_i^*} = \frac{1}{\gamma_{ij}}\left(1 - \mu_{ij}\right). \tag{S16}$$

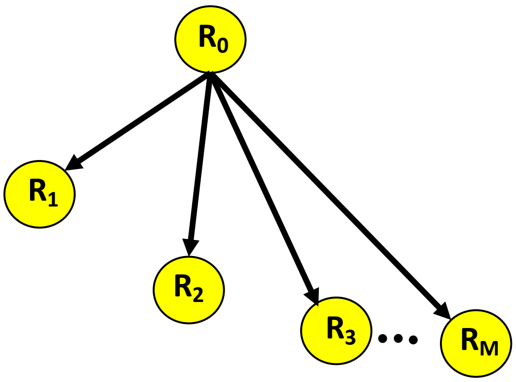


Figure S1: **The reactions in the hub and spokes network.** All reactions start from $R_0$.

In addition to the above, the steady state equations also give us an additional condition:

$$R_0^* = h_0 - \sum_i R_i^*. \tag{S17}$$

This is equivalent to flux conservation, the total inflow of resources must match the total outflow of resources.

Together, we can derive the steady-state concentration of all the resources. We will explain the solution for a general network after illustrating the solution for two special limiting network scenarios.

#### Hub and spokes network

The first network we consider is a hub and spoke network (Fig. S1). All active reactions emanate from the sole supplied resource $R_0$, giving products $R_1..R_M$. We can solve the system of equations we derived above for this network in a straightforward manner. Eq.(S16) reduces to:

$$\frac{R_i^*}{R_0^*} = \frac{1}{\gamma_{0i}} \left(1 - \mu_{0i}\right) \;\; \forall i \in \{1...M\}. \tag{S18}$$

Solving these equations and Eq.(S17) we get:

$$R_0^* = \frac{h_0}{1 + \sum_{j=1}^{M} \frac{1}{\gamma_{0j}} \left(1 - \mu_{0j}\right)}, \tag{S19}$$

and

$$R_i^* = R_0^* \frac{1}{\gamma_{0i}} \left(1 - \mu_{0i}\right). \tag{S20}$$

#### Linear chain network

The second network we consider is the opposite limit, of a linear reaction chain (Fig. S2). The active reactions connect resources closest in energy. Thus the network forms a chain starting at $R_0$, proceeding down the resource energy hierarchy, and ending at the lowest energy resource $R_M$. Here, Eq.(S16) reduces to:

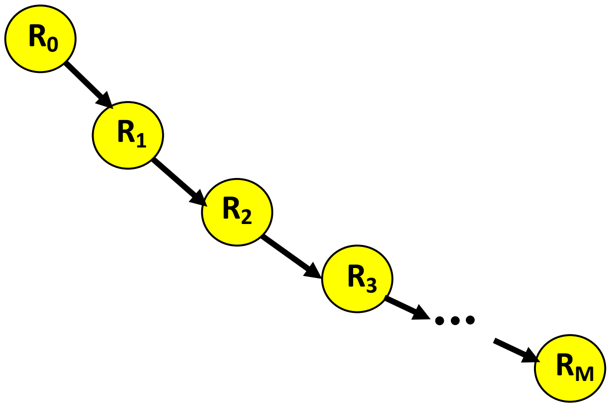


Figure S2: **The linear reaction chain network.**

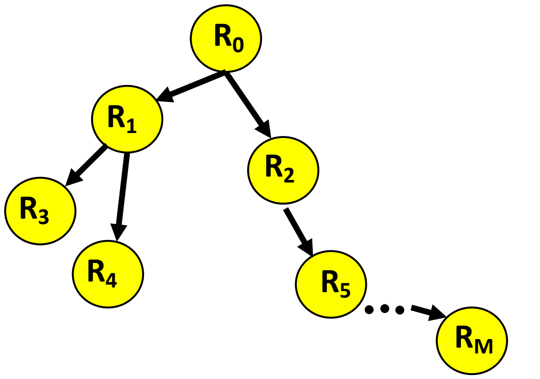


Figure S3: **A general tree network.** Resource nodes can have any number of outflow reactions to lower energy products. No node can have more than one inflow reaction.

$$\frac{R_i^*}{R_{i-1}^*} = \frac{1}{\gamma_{i-1,i}} \left(1 - \mu_{i-1,i}\right) \quad \forall i \in \{1...M\}. \tag{S21}$$

We can recursively solve these equations to express every resource concentration in terms of $R_0^*$, and use Eq.(S17) to get:

$$R_0^* = \frac{h_0}{1 + \sum_{j=1}^{M} \prod_{k=1}^{j} \frac{1}{\gamma_{k-1,k}} \left(1 - \mu_{k-1,k}\right)}, \tag{S22}$$

and

$$R_i^* = R_0^* \prod_{k=1}^{i} \frac{1}{\gamma_{k-1,k}} \mu_{k-1,k}. \tag{S23}$$

This expression clarifies that when reactions occur serially, we have a separate product appear for each resource in the chain.

#### General network

In the slow-growth limit, the in-degree of a resource is constrained to be 1—thus the network is a tree. We can solve the steady-state equations for an arbitrary tree network using the insights gained from solving the special limits above.

We have the following steady state equations for the tree network:

$$\frac{R_j^*}{R_i^*} = \frac{1}{\gamma_{ij}}\left(1-\mu_{ij}\right) \;\; \forall (i,j) \in \mathcal{X}, \tag{S24}$$

where $\mathcal{X}$ is the set of active reactions in the network. To solve this system, we need to express the steady-state concentration of each resource in terms of the supplied resource, $R_0$. Since the network is a tree, $R_0$ is connected to a resource $R_j$ via a unique path, composed of multiple reactions. We represent the path in short as $R_0 \to R_j$, and the set of reaction on the path as $\mathcal{P}_{0j}$. Each path resembles the linear chain solved in the previous section. Using this insight, we find the steady-state concentrations as:

$$R_0^* = \frac{h_0}{1+\sum_{k=1}^{M}\prod_{i,j\in\mathcal{P}_{0\to k}}\frac{1}{\gamma_{ij}}\left(1-\mu_{ij}\right)}, \tag{S25}$$

The pathway to each possible product contributes a term to the sum. To leading order, this term is a product of the free energy dissipated on the pathway.

The steady state solution for $R_k$ is

$$R_k^* = R_0^* \prod_{i,j\in\mathcal{P}_{0k}} \frac{1}{\gamma_{ij}}\left(1-\mu_{ij}\right) \tag{S26}$$

When the steady-state growth of a community is very slow ($\mu_{ij} \ll 1$ ), we have

$$R_k^* \approx R_0^* \prod_{i,j\in\mathcal{P}_{0k}} \frac{1}{\gamma_{ij}} \tag{S27}$$

Note that the above analysis assumes that the identities of the surviving species and active reactions are provided. It does not consider competition between species and explain which reactions will be active in the community. We will consider species competition in the following section.

### S3 Selection of metabolic network: principle of maximum dissipation

In the preceding section, we solved for the steady-state community assuming that we know the active reactions and the surviving species that utilize them. In this section, we will identify the reactions that survive in the community.

From an ecological point of view, we can perform an invasion analysis to identify the active reactions. Given a steady state community, we can consider the invasion of species $\beta_{ik}$ specializing on a reaction $R_i \to R_k$ that is inactive in the current community (Fig. S4). For the invader to succeed, we need it to have a positive growth rate in the environment it is invading, i.e.,

$$k_{cat,ik}E_{ik}^{(\beta_{ik})}Y\mathcal{E}_{ik}^{\mathrm{ATP}}\left(1-\gamma_{ik}\frac{R_k^*}{R_i^*}\right)-\delta > 0, \tag{S28}$$

where $R_k^*, R_i^*$ are the steady-state resource concentrations in the community prior to invasion. Using (Eq.(S26)), we can simplify this invasion condition as

$$\frac{\prod_{u,v\in\mathcal{P}_{0k}}\frac{1}{\gamma_{uv}}\left(1-\mu_{uv}\right)}{\prod_{u\prime,v\prime\in\mathcal{P}_{0i}}\frac{1}{\gamma_{u\prime v\prime}}\left(1-\mu_{u\prime v\prime}\right)} < \frac{1}{\gamma_{ik}}\left(1-\mu_{ik}\right), \tag{S29}$$

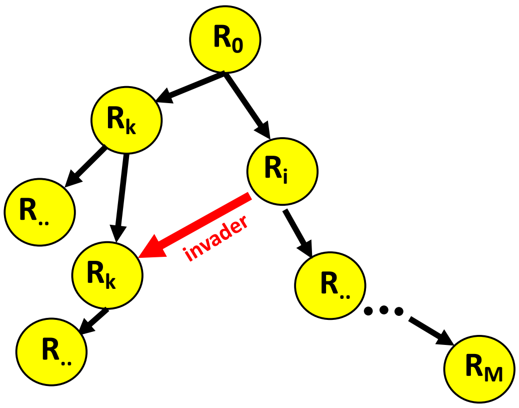


Figure S4: **Analysing invasion of an alternative path reveals network selection principle.** A steady-state metabolic network, depicted by the black arrows, is invaded by a species utilizing the red path from $R_i$ to $R_k$. This path could be composed of multiple reactions. The conditions determining the survival of the invader shows that the network is selected by the maximum dissipation principle.

where the paths $\mathcal{P}_{0k}, \mathcal{P}_{0i}$ refer to the reaction paths in the community before invasion

We are interested in the slow-growth limit, which is equivalent to small dilution rate $\delta$. For small enough $\delta$ such that $\mu_{ij} = \frac{\delta}{k_{cat,ij}\mathcal{E}_{ij}^{\mathrm{ATP}} Y E(\beta)_{ij}} \ll 1$ for any reaction, the above expression simplifies to

$$\frac{\prod_{u,v\in\mathcal{P}_{0k}} \frac{1}{\gamma_{uv}}}{\prod_{u\prime,v\prime\in\mathcal{P}_{0i}} \frac{1}{\gamma_{u\prime v\prime}}} < \frac{1}{\gamma_{ik}} \tag{S30}$$

Using the definition of $\gamma$, this implies that the invader successfully invades if

$$-\Delta G^0_{0\to k} < -\Delta G^0_{0\to i} - \Delta G^0_{ik}, \tag{S31}$$

where $-\Delta G^0_{0\to k}$ and $-\Delta G^0_{0\to i}$ refer to the total free energy dissipated over the reactions on the paths $R_0 \to R_k$ and $R_0 \to R_i$ in the steady state community before invasion, and $-\Delta G^0_{ik}$ is the free energy dissipated in the reaction utilized by the invader. Hence the RHS equals the free energy dissipated on the new path from $R_0$ to $R_k$ created by the invader, and the equation is comparing the free energy dissipated on the two possible paths to $R_k$. Thus, the invader is able to successfully invade only if the new path to $R_k$ dissipates more free energy than the path to $R_k$ in the resident community.

Over many invasions, the community metabolic network will converge to utilize the paths to each resource that dissipates most free energy, and therefore be uninvasible. Thus the principle of maximum dissipation determines the reaction network that is selected by the final community.

Note that the final community can still be successfully invaded by species utilizing the maximally dissipative reactions but happen to be better competitors due to either allocation more enzyme to these reactions or having better enzymes to utilize these reactions. These effects are described in the $\mu$ dependence of Eq.(S29), which appears as a lower order contribution for slow-growing communities. Thus communities could continuously experience species turnover while maintaining their metabolic functions.

An alternate way to understand the principle of maximum dissipation is to consider the ratio of $R_k^*/R_0^*$ in the communities differing only by the reaction leading to $R_k$. The community dissipating more free energy on the path to $R_k$ sustains a higher value of $R_k^*/R_0^*$. Thus the species utilizing this maximally dissipative path is able to invade the second community successfully but not vice-versa. Since this argument applies to all resources $R_1 ... R_M$, the final community, that has survived many invasions, utilizes the maximally dissipative path.

### S4 Convergence of metabolic fluxes

The fluxes in the active reactions in the community can be calculated using our results for the steady-state resource concentrations (Eqs.(S25),(S26)) by applying one additional physical principle: conservation. The resource flux via reactions into each resource node in the network is balanced by the flux out of the node via reactions and dilution. The dilution flux of a resource is determined by the dilution rate. Thus, the total flux into resource $R_i$, $I_i$, is (also Eq.7 in the main text)

$$I_i^* = \delta R_i^* + \sum_{j \text{ downstream of } i} \delta R_j^*, \tag{S32}$$

where $j$ runs over all resources that appear as products in reactions downstream of $R_i$. Since the resource concentrations are set by thermodynamics (to leading order), the fluxes in the community also converge to this thermodynamically determined value. Thus we observe functional convergence in the community.

### S5 Incorporating maintenance energy requirements

We can incorporate the maintenance energy requirements of a cell by replacing $\delta$ in Eq. (S7) such that

$$\frac{dN_\alpha}{dt} = N_\alpha \left[ \sum_{i,j \in \mathcal{X}} E_{ij}^{(\alpha)} \mathcal{F}_{ij} \mathcal{E}_{ij}^{\text{ATP}} Y - \delta - m_\alpha \right], \tag{S33}$$

where $m_\alpha$ is the energy flux required for cell maintenance. If the variation in maintenance costs between species is small compared to the average maintenance cost, $m$, the analysis of the model without maintenance costs can be repeated in a straightforward manner.

The equations describing resource concentrations all apply with a new definition of $\mu_{ij}$, given by

$$\mu_{ij} = \frac{\delta + m}{k_{cat,ij} \mathcal{E}_{ij}^{\text{ATP}} Y} \left[ \mathbf{E^{-1}}.\vec{1} \right]_{ij}, \tag{S34}$$

where terms capturing the species-dependent variation in maintenance energy are neglected as small corrections to $\mu_{ij}$. Once again, we find that resource concentrations are determined by thermodynamics, to leading order in $(\delta + m)/g_{max}$. From Eq. (S29), we see that the maximum dissipation principle determines the network selected by ecological competition.

Again, due to flux balance we can use Eq. (S32) to calculate the fluxes in each reaction. Note that the reaction flux is still proportional to $\delta$ because resource loss in this model only occurs via dilution. This resource dilution is essential to ensure that the community is at a non-equilibrium steady-state.

### S6 Generalising to arbitrary reaction stoichiometry

In this section we generalize the model to the scenario where the stoichiometry of the substrate to product reaction is no longer $1 : 1$. We will show that our main results of maximum dissipation

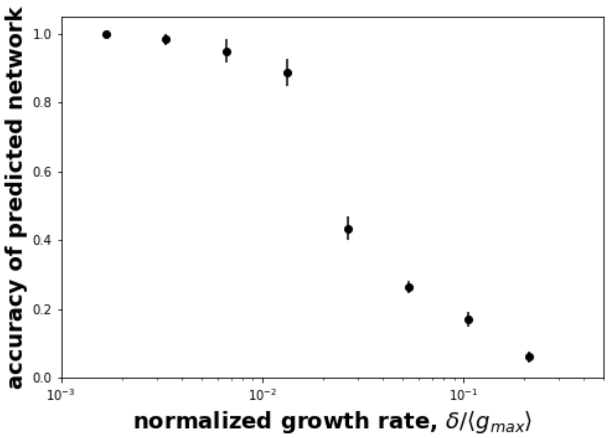


Figure S5: **Maximum dissipation predicts metabolic network accurately for larger variation in enzymatic parameters.** The accuracy of the network predicted by maximum dissipation was quantified as the fraction of reactions in community metabolic network correctly predicted. The enzymatic parameters were picked from lognormal distributions with $\mu = 1$ and $\sigma = 0.15$. Results were average over ten species pools with enzyme budget and allocation chosen similarly as in Fig.2 in the main text. The environment was the same as in Fig.2 in the main text.

and functional convergence are robust to changes in stoichiometry. Stoichiometry only modifies the equations quantitatively.

Specifically, we consider the enzyme catalyzed reversible reaction

$$E + y_s S \rightleftharpoons E + y_p P, \tag{S35}$$

where $y_s, y_p$ are the stoichiometric coefficient of the substrate and product in the reaction. The free energy dissipated in the reaction, defined as

$$-\Delta G^0_{SP} = y_s \mathcal{E}^0_S - y_p \mathcal{E}^0_P - \mathcal{E}^{\mathrm{ATP}}, \tag{S36}$$

accounts for reaction stoichiometry. The reaction flux is again described by reversible Michaelis-Menten kinetics [7, 6]:

$$J = k_{cat} E \frac{\left(\frac{S}{K_S}\right)^{y_s}}{1 + \left(\frac{S}{K_S}\right)^{y_s} + \left(\frac{P}{K_P}\right)^{y_p}} \left(1 - \frac{P^{y_p}}{S^{y_s}} \gamma\right), \tag{S37}$$

where $\gamma = e^{\Delta G^0_{SP}/RT}$ and the flux per unit enzyme being $F = J/E$.

We now consider a full connected network of resources $R_0, R_1, ...R_M$, ordered in terms of their standard-state energies as before. The stoichiometric coefficients of the reaction in the network cannot be entirely arbitrary. For e.g., a mole of glucose (with 6 carbons), can produce no more than 3 moles of acetate (with 2 carbons). Motivated by this, for our reaction network, we enforce the constraint that the number of moles of resource $R_i$ produced from a single mole of $R_0$ should be independent of the reaction path. As a concrete example, the two reaction paths from $R_0$ to $R_2$ will have the reactions

$$y_0 R_0 \xrightleftharpoons{E_{01}} y_1 R_1, \xrightleftharpoons{E_{12}} y_2 R_2,$$

$$y_0 R_0 \xrightleftharpoons{E_{02}} y_2 R_2.$$

Note that the stoichiometric coefficient of $R_2$ in both reactions is the same, $y_2$, due to this stoichiometric constraint. Without loss in generality, we assume that the stoichiometric coefficient of $R_0$, $y_0$ is equal to 1. This allows us to interpret the stoichiometric coefficient of each resource in the network as the number of moles produced per mole of the supplied resource $R_0$.

The equations for species and resource dynamics extend in a straightfoward manner as:

$$\frac{dN_\alpha}{dt} = N_\alpha \left[ \sum_{i,j \in \mathcal{X}} E_{ij}^{(\alpha)} \mathcal{F}_{ij} \mathcal{E}_{ij}^{\text{ATP}} Y - \delta \right], \tag{S38}$$

and

$$\frac{dR_i}{dt} = h(R_i) - \sum_{\alpha,j} y_i N_\alpha E_{ij}^{(\alpha)} \mathcal{F}_{ij} + \sum_{\alpha,k} y_i N_\alpha E_{ki}^{(\alpha)} \mathcal{F}_{ki} - \delta R_i. \tag{S39}$$

We can again solve for the steady-state by using the inverse of the enzyme allocation matrix of the survivors as above. The analog of Eq. (S12) will now be

$$\left( 1 - \gamma_{ij} \frac{\left(R_j^*\right)^{y_j}}{\left(R_i^*\right)^{y_i}} \right) = \frac{\delta}{k_{cat,ij} \mathcal{E}_{ij}^{\text{ATP}} Y} \frac{1 + \left(\frac{R_i^*}{K_{S_{ij}}}\right)^{y_i} + \left(\frac{R_j^*}{K_{P_{ij}}}\right)^{y_j}}{\left(\frac{R_i^*}{K_{S_{ij}}}\right)^{y_i}} \left[ \mathbf{E^{-1}} \vec{1} \right]_{ij}, \tag{S40}$$

Like above, we can simplify this in the high-flux limit (Eq. (S6)), where reaction fluxes are primarily slowed down by thermodynamic inhibition. For each active reaction in the final community, we get the analog of Eq. (S13) :

$$\frac{\left(R_j^*\right)^{y_j}}{\left(R_i^*\right)^{y_i}} = \frac{1}{\gamma_{ij}} \left( 1 - \frac{\delta}{k_{cat,ij} \mathcal{E}_{ij}^{\text{ATP}} Y} \left[ \mathbf{E^{-1}} \vec{1} \right]_{ij} \right). \tag{S41}$$

Thus $\frac{\left(R_j^*\right)^{y_j}}{\left(R_i^*\right)^{y_i}}$ is set by thermodynamics to leading order in $\delta$.

To simplify the notation, we will define $\mu_{ij}$ once again as

$$\mu_{ij} = \frac{\delta}{k_{cat,ij} \mathcal{E}_{ij}^{\text{ATP}} Y} \left[ \mathbf{E^{-1}} . \vec{1} \right]_{ij} \tag{S42}$$

We can use the same procedure as above to express the concentration ratio of two resources for an arbitrary tree-like metabolic network in terms of the free energy dissipated on the path between the two resources.

$$\frac{\left(R_j^*\right)^{y_j}}{\left(R_i^*\right)^{y_i}} = \prod_{u,v \in \mathcal{P}_{ij}} \frac{1}{\gamma_{uv}} \left(1 - \mu_{uv}\right), \tag{S43}$$

where $\mathcal{P}_{ij}$ is set of reactions on the path from $R_i$ to $R_j$. To leading order in $\delta$ we get :

$$\frac{\left(R_j^*\right)^{y_j}}{\left(R_i^*\right)^{y_i}} = e^{-\Delta G^0_{i \Rightarrow j}/RT} + \mathcal{O}\left(\frac{\delta}{g_{\max}}\right). \tag{S44}$$

This equation shows that the ratio of steady-state resource concentrations (raised to powers based on stoichiometric coefficients) on an arbitrary tree network is once again determined by the free energy dissipated on the path between the two resources.

We now study the question of which steady-state network gets selected as a result of ecological competition between species. Specifically, we consider the outcome of competition between species using different paths to the same resource, $R_k$. Like before, for a given steady state community, we consider the invasion of species $\beta_{ik}$ specializing on a reaction $R_i \to R_k$ that is inactive in the current community (Fig. S4). For the invader to succeed, we need it to have a positive growth rate in the environment it is invading, i.e.,

$$k_{cat,ik} E_{ik}^{(\beta_{ik})} Y \mathcal{E}_{ik}^{\mathrm{ATP}} \left(1 - \gamma_{ik} \frac{(R_k^*)^{y_k}}{(R_i^*)^{y_i}}\right) - \delta > 0, \tag{S45}$$

where $R_k^*, R_i^*$ are the steady-state resource concentrations in the community prior to invasion. Using (Eq.(S44)), we can simplify this invasion condition as

$$\frac{\prod_{u,v \in \mathcal{P}_{0k}} \frac{1}{\gamma_{uv}} (1 - \mu_{uv})}{\prod_{u\prime,v\prime \in \mathcal{P}_{0i}} \frac{1}{\gamma_{u\prime v\prime}} (1 - \mu_{u\prime v\prime})} < \frac{1}{\gamma_{ik}} (1 - \mu_{ik}), \tag{S46}$$

where the paths $\mathcal{P}_{0k}, \mathcal{P}_{0i}$ refer to the reaction paths in the community before invasion. For small enough $\delta$ such that $\mu_{ij} = \frac{\delta}{k_{cat,ij} \mathcal{E}_{ij}^{\mathrm{ATP}} Y E^{(\beta)}_{ij}} \ll 1$ for any reaction, the above expression simplifies to

$$-\Delta G^0_{0\to k} < -\Delta G^0_{0\to i} - \Delta G^0_{ik}, \tag{S47}$$

we have used the definition of $\gamma$. $-\Delta G^0_{0\to k}$ and $-\Delta G^0_{0\to i}$ refer to the total free energy dissipated over the reactions on the paths $R_0 \to R_k$ and $R_0 \to R_i$ in the steady state community before invasion, and $-\Delta G^0_{ik}$ is the free energy dissipated in the reaction utilized by the invader. Hence the RHS equals the free energy dissipated on the new path from $R_0$ to $R_k$ created by the invader. Thus the invader is able to successfully invade only if the new path to $R_k$ dissipates more free energy than the path to $R_k$ in the resident community. So once again we discover that the principle of maximum dissipation determines the reaction network selected by ecological competition of many species.

We now address the question of functional convergence. Flux conservation in the reaction network gives us

$$\sum_{i=0}^{M} y_i R_i^* \delta = h_0 \delta, \tag{S48}$$

where $h_0$ is the concentration of the resource supply and $y_0 = 1$. Using Eq. (S44), we can write this in terms of only $R_0^*$:

$$\sum_{i=0}^{M} y_i \left(\frac{R_0^*}{\gamma_{0i}}\right)^{1/y_i} - h_0 = 0. \tag{S49}$$

Unlike the 1 : 1 stoichiometry scenario, here we get a nonlinear equation for $R_0$. A nonlinear equation could have multiple solutions for $R_0^*$, which would correspond to alternative steady-states of community function. However, because the stoichiometric coefficients $y_i$ are all positive we are guaranteed to have only a single positive solution for $R_0^*$ by Descartes rule of signs (extended to rational exponents) [8]. We can also understand this intuitively by noticing that the function is monotonically increasing for positive $R_0^*$. Thus we have a single positive solution for $R_0^*$, which we can obtain numerically.

Using the solution for $R_0^*$,Eq. (S44), and flux conservation at each resource node, we can calculate the fluxes in each reaction as given by Eq. (S32). Thus we have functional convergence for arbitrary reaction stoichiometries.

### S7 A model without resource dilution

We now consider a model without resource dilution. Without any flow of resource out of the system, the system cannot reach a non-trivial steady-state—the constant inflow of resources will eventually clog up the system. Here, we show that an effective outflow of resources can appear from anabolism accompanied by species death. Again, the principle of maximum dissipation determines the community metabolic network.

In this model, resources are used to maintain biomass through anabolic processes by species at resource-specific rates $b_i$. A species $\alpha$ dies or sinks with rate $m_\alpha$ . The accompanying loss of species biomass creates a flux of resources out of the system, which allows the system to attain a non-equilibrium steady-state.

The equation describing dynamics of species $\alpha$ with abundance $N_\alpha$ is

$$\frac{dN_\alpha}{dt} = N_\alpha \left[ \sum_{i,j} E_{ij}^{(\alpha)} \mathcal{F}_{ij} Y - m_\alpha \right], \tag{S50}$$

where $\mathcal{F}$ denotes the per-capita flux as defined previously.

The dynamics of resource $i$ with concentration $R_i$ is given by

$$\frac{dR_i}{dt} = h(R_i) + \sum_{\alpha,k} N_\alpha E_{ki}^{(\alpha)} \mathcal{F}_{ki} - \sum_{\alpha,j} N_\alpha E_{ij}^{(\alpha)} \mathcal{F}_{ij} - \sum_{\alpha} N_\alpha b_i R_i = 0, \tag{S51}$$

where the first and second terms account for catabolic reactions where $R_i$ acts as the product and substrate respectively, and the last term accounts for resource-assimilation for biomass (anabolism) and maintenance at resource-dependent rates $b_i$. Since resource-consumption in anabolism and maintenance is driven by ATP (energy assimilated by the microbe), we do not consider thermodynamic limitation in this term. Hence, even the lowest energy resource can be consumed by the microbe. We assume that $b_i$ is very small and the resource needs for the slow growth and maintenance will be small.

Again, we assume only a single resource, $R_0$ is supplied with $h(R_0) = h_0/\tau$, with $\tau$ setting the time scale of supply to allow $h_0$ to retain the dimensions of resource concentration as before.

For convenience and clarity, we study a system of specialist species . At steady-state, species growth needs to balance loss due to maintenance and/or sinking. Therefore, we have

$$\frac{R_i^*}{R_j*} = \frac{1}{\gamma_{ij}} \left( 1 - \frac{m_\beta}{k_{cat,ij} \mathcal{E}_{ij}^{\mathrm{ATP}} Y E_{ij}^{(\beta)}} \right), \tag{S52}$$

for every active reaction in the community. $\beta$ is the specialist species catalyzing reaction between $R_i$ and $R_j$.

For slow-growing communities, $m_\beta$ is small. Therefore, the ratio $\frac{R_i^*}{R_j*}$ depends only on the free energy dissipated in the corresponding reaction, $\gamma_{ij}$, to leading order.

Repeating the analysis applied to the previous model, we find again that the ratio $\frac{R_i^*}{R_0*}$ depends only the free energy dissipated along the realized reaction path from $R_0$ to $R_i$. Similarly, we also find that the principle of maximal dissipation determines the set of realized reactions in the network.

To arrive at the steady state concentration of resource, we use flux conservation, which reads as

$$\frac{h_0}{\tau} - \sum_{\alpha,i} N_\alpha^* b_i R_i^* = 0, \tag{S53}$$

or rather,

$$R_0^* = \frac{h_0}{\tau b_0 N_{\text{tot}}^*} - \sum_{i=1}^{M} \frac{b_i}{b_0} R_i^*, \tag{S54}$$

where $N_{\text{tot}}^* = \sum_\alpha N_\alpha^*$ is the total biomass in the community. Resource-consumption by species for anabolism and maintenance, and the subsequent death and sinking of the species creates an out-flux of resources that balances the influx of $R_0$ and creates a steady-state.

And now, we can calculate the steady-state solution for resource concentrations:

$$R_0^* = \frac{\frac{h_0}{\tau b_0 N_{\text{tot}}^*}}{1 + \sum_{i=1}^{M} \frac{b_i}{b_0} e^{-\Delta G^0_{0\Rightarrow i}/RT}}, \tag{S55}$$

and

$$R_i^* = R_0^* e^{-\Delta G^0_{0\Rightarrow i}/RT}, \tag{S56}$$

in the limit of slow growth, where $e^{-\Delta G_{0\Rightarrow i}}$ is the free energy dissipated along the path from $R_0$ to $R_i$. Note that these solutions are not closed form; the solution depends on the total species biomass, $N_{\text{tot}}^* = \sum_\alpha N_\alpha^*$. However, as long as the total biomass of communities assembled from different species pools in replicate environments are similar, we expect the resource concentrations in the environments to be the same. Also note that if $b_i$ is large, then substrate depletion can be a major form of competition between species, and our assumption that reactions are slowed primarily by thermodynamic inhibition does not hold.

The net flux into a resource $R_i$ is then given by

$$I_i^* = N_{\text{tot}}^* b_i R_i^* + \sum_{j \text{ downstream of } i} N_{\text{tot}}^* b_j R_j^* \tag{S57}$$

Thus we have approximate functional convergence as well, with a new expression for the reaction fluxes. The functional convergence is now sensitive to variation in the total species abundance, across replicates.

### S8 Modifying the energy dissipation of individual reactions in the network

In these simulations, we studied the effect of changing the free energy dissipated in a single reaction to a large value $\geq 10RT$. This represents a reaction in the network become almost irreversible while other reactions are reversible. The resource energies were large to allow reactions to become irreversible while remaining feasible under standard conditions. Specifically, the energies of the resources $R_0, R_1, R_2, R_3, R_4$, and $R_5$ were $60RT, 48RT, 36RT, 24RT, 12RT, 0RT$ in Fig. S7A and $80RT, 64RT, 48RT, 32RT, 16RT, 0RT$ in Fig. S7B .

To ensure that all reactions except the modified reaction were reversible, we randomly sampled the energy dissipated in the reaction using an assumption motivated by quantization of energy assimilation [9, 10, 11]. Specifically, we assumed that when cells try to assimilate as much energy as possible, the energy dissipated in each reaction will be less than one quantum of energy (chosen $= 5RT$). Then, we picked a separate uniform random number between 0 and $5RT$ as the energy dissipated for each reaction. We repeated this random sampling to get 50 different realizations of the chemical networks or "worlds" with different values of energy dissipation. In each world, we simulated 5 different species pools to estimate the amount of functional convergence in the world.

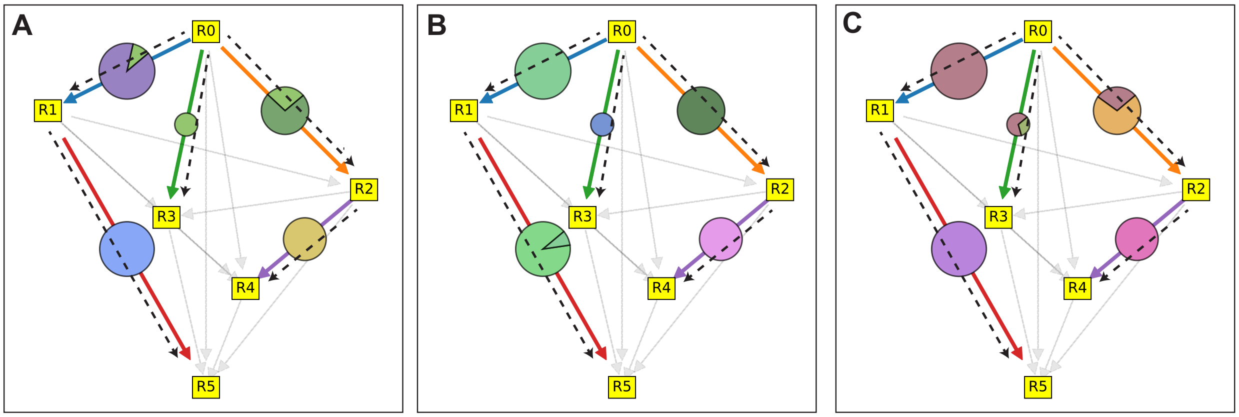


Figure S6: **Maximum dissipation principle and functional convergence in model without resource dilution.** The panels show the metabolic network of three communities assembled from three different species pools in identical environments. The maximum dissipation principle predictions, shown by dashed lines, predicts the reactions accurately. The fluxes through the reactions, represented by the size of the circles on the corresponding arrows, are similar across the three communities. The pie-charts depict how the flux is shared between species in the different communities. This differs between communities. Parameters for the figure were: $b_i$ was chosen from as $10^{-}3$ times a lognormal random variable with parameters $\mu = 0, \sigma = 0.01$. The maintenance cost was $m = 0.01$, resource supply was described by $h_0 = 80$ and $\tau = 10$. Resource energies are the same as in Fig.2 in the main text. The energy assimilated and enzymatic parameters are chosen from the same distributions as in Fig.2 in the main text.

Thus, we simulated a total of 250 pools across 50 different worlds. The concentration of $R_0$ in the supply, $h_0 = 80$. The chemical kinetic parameters $K_P$,$K_S$, and $k_{cat}$ were chosen from lognormal distributions with lognormal parameters $\mu = 0, \sigma = 0.01$ for each pool. The energy to biomass yield factor, $Y = 1$. The dilution rate $\delta = 0.01$.

For each pool, we ran 15 separate experiments which modified the energy dissipated in each of the 15 separate reactions, one at a time, to a large value of $10RT$ in Fig. S7A and $12RT$ in Fig. S7B. In each world, we computed the average accuracy of the predicted metabolic network and the functional distance between the 5 different pools as a function of the reaction modified. We report these values averaged over the 50 different worlds. Irrespective of the reaction modified, we found that the accuracy of the network predicted was greater than expected by chance (Fig. S7). The expected accuracy if the reaction network realized in the community was a random maximum spanning tree was 0.31 (dashed line in the figure). Thus, our theoretical predictions retained some utility despite the high energies in the network.

We also investigated the effect of the location of the modified reaction. Specifically, when we ordered the modified reactions by their average position, we found agreement with theory and functional convergence was higher if the modified reactions were located near the top of the network. In other words, having more irreversible reactions near the top of the metabolic network was less disruptive. In Fig. S7 the reactions are ordered by average position in the network defined as the average of the substrate index and product index; the reaction $R0 \to R2$ thus had an average position of 1. In case two reactions were tied, such as $R0 \to R4$ and $R1 \to R3$, we assumed the reaction with the substrate nearer to the top of the network had a lower average position. Using this definition of ordering reactions, we found a significant ($p < 0.05$) rank correlation $\rho$ between average accuracy of the predicted metabolic network and average position of the reaction as well as

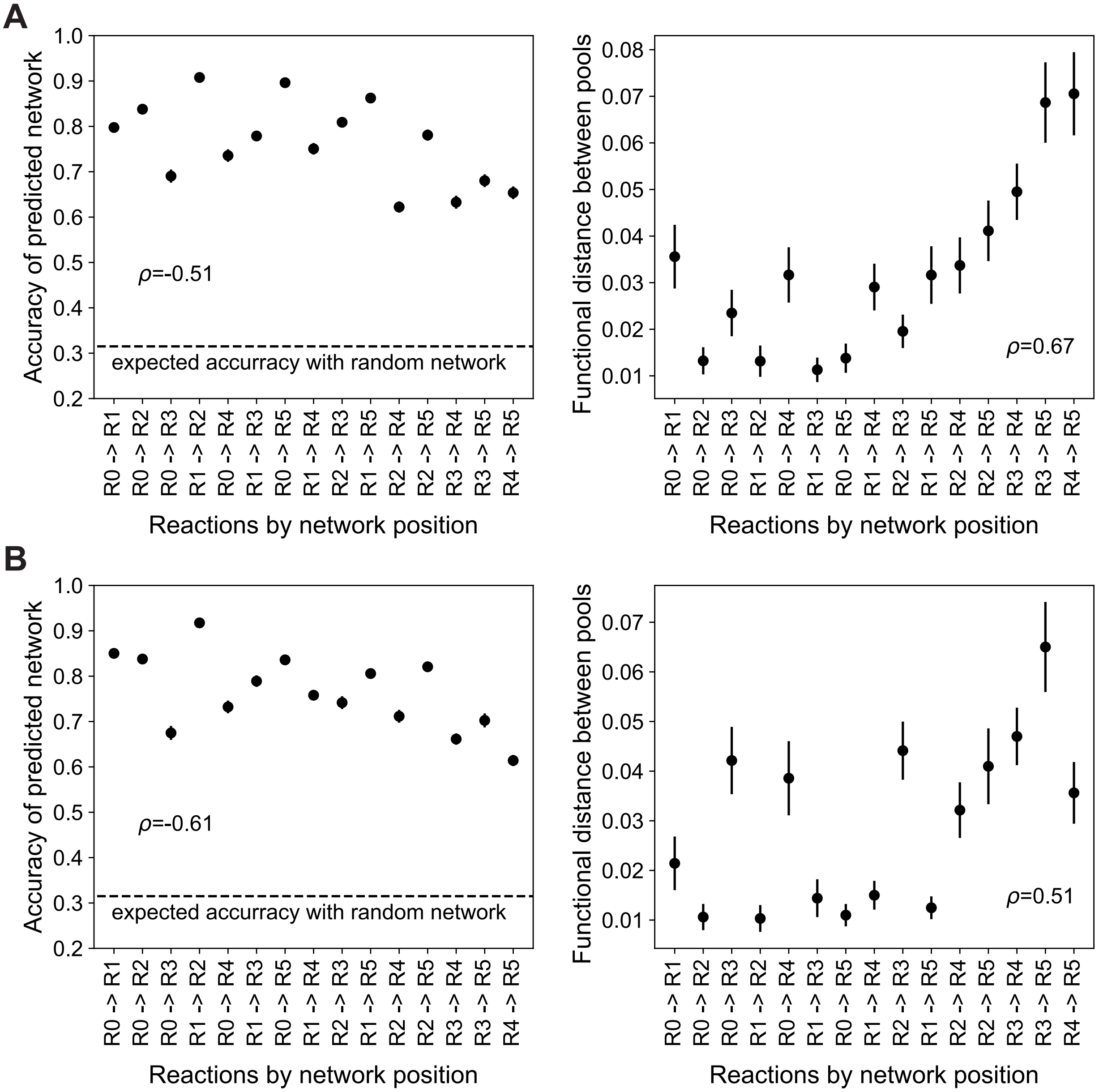


Figure S7: **Highly dissipative reactions near the top of the metabolic network are less disruptive.** **A)** The accuracy of the network predicted by maximum dissipation decreases as reactions towards the bottom of the network are made more irreversible. The dashed line indicates the expected accuracy if the observed network was a random spanning tree. The functional distance between fluxes from different pools increases as reactions towards the bottom of the network are made more irreversible. $\rho$ indicates the spearman rank correlation. **B)** Same trend is observed in a simulations with different parameters. All reported correlations were significantly $p < 0.05$ different from zero.

average functional distance and average position of the reaction. These correlations indicate that modifying reactions near the top of the network was less disruptive. This is potentially more likely to occur in real environments where the resources at the top of the metabolic network are typically of higher energy while reactions towards the bottom of the network are lower in energy.

#### S9 How much free energy dissipation makes a reaction irreversible?

Reversible Michaelis-Menten kinetics (S3) describes the simplified catabolic reaction we consider for all ranges of free energy dissipation. But from physical considerations we expect that for large enough values of free energy dissipation, the thermodynamic product inhibition term $\left(1 - \frac{P}{S} e^{\Delta G^0/RT}\right)$ will become irrelevant. But how large is large enough?

To answer this question in a self-consistent manner, we estimated the ratio $\frac{P}{S}$ by artificially ignoring thermodynamic inhibition. Specifically, we considered a species growing using an irreversible catabolic reaction converting substrate $S$ to product $P$. Species dynamics is described by the irreversible equivalent of Eq. (S7) in this single reaction scenario:

$$\frac{dN}{dt} = N\left[k_{cat} E \frac{S}{K_S} \mathcal{E}^{\mathrm{ATP}} Y - \delta\right], \tag{S58}$$

where the parameters are as described before.

At steady state, this gives us

$$S^* = \frac{\delta K_S}{E k_{cat} \mathcal{E}^{\mathrm{ATP}} Y}. \tag{S59}$$

Thus due to the lack of product inhibition, the substrate is depleted down to a level that is proportional to the dilution rate $\delta$. The substrate concentration in the supply is $h_0$ and hence the supply flux of the substrate is $h_0\delta$. Due to flux conservation, we have $h_0\delta = S^*\delta + P^*\delta$. Since we are working in the the regime where concentration in the supply is large and dilution rate is small, this implies $P^* \approx h_0$. Together this gives us the steady state ratio of $\frac{P^*}{S^*}$ as

$$\frac{P^*}{S^*} = \frac{h_0 E k_{cat} \mathcal{E}^{\mathrm{ATP}} Y}{\delta K_S}. \tag{S60}$$

The above equation provides the concentration ratio that would be maintained by a single species on a single reaction if there was no thermodynamic inhibition.

For the substrate to be depleted down to the level specified by the above equation, the thermodynamic inhibition should be negligible. In other words, $\left(1 - \frac{P^*}{S^*} e^{\Delta G^0/RT}\right) \approx 1$ or

$$e^{\Delta G^0/RT} \ll \frac{\delta K_S}{h_0 E k_{cat} \mathcal{E}^{\mathrm{ATP}} Y}. \tag{S61}$$

This equation gives us the free energy dissipation required for thermodynamic inhibition to be irrelevant and reaction to be effectively irreversible. If we know the parameter values on the right hand side of the equation, we can estimate the free energy dissipation required for making a reaction effectively irreversible.

We obtained estimates of these parameters from the literature to estimate the free energy scale that would make a reaction irreversible. The parameters values vary between species, metabolic pathways, enzymes and experimental conditions. Therefore, we used appropriate, representative values measured to obtain a rough estimate of the free energy scale that would make a reaction irreversible.

We obtained the dilution rate $\delta$ and supply concentration $h_0$ from the anaerobic digester experiments we analyzed in this study [12]. A dilution time of 15 days implies $\delta = 7.7 \times 10^{-7}\text{sec}^{-1}$. The influent concentration in the study was $15\text{gCOD} \cdot \text{L}^{-1}$, where gCOD is denotes the number of grams of oxygen required to completely oxidize the substrate (Chemical Oxygen Demand). We convert this into the number of electrons the influent can donate since the substrate is the electron donor. Since one mole of oxygen gas weighs 32g and accepts four electrons, the influent concentration of $15\text{gCOD} \cdot \text{L}^{-1}$ becomes $\approx 2\,\text{mol-electron} \cdot \text{L}^{-1}$ in redox terms.

The enzyme kinetic parameters are estimated from Refs. [13, 14]. (Note that $K_S$ is a Michaelis constant usually denoted as $K_M$.) The ratio $k_{cat}/K_S$ has a median value of $4 \times 10^5\text{M}^{-1} \cdot \text{sec}^{-1}$ in central carbon metabolism. Since we are considering the major catabolic pathway of the cell, we use an enzyme copy number of $10^4$ per cell which estimates the copy number of highly expressed enzymes in the cell [14]; this is equivalent to $1.66 \times 10^{-20}$ moles per cell. Combined with the substrate concentration in the supply, we get a chemical flux of the moles of electrons produced per second in catabolism as $\frac{h_0 k_{cat} E}{K_S} = 1.33 \times 10^{-14}\,\text{mol-electron} \cdot \text{sec}^{-1}$.

We now need estimates for $\mathcal{E}^{\text{ATP}}Y$, which converts the chemical concentration flux to yield in terms of number of cells. Roden and Jin [15, 16] estimates 10.5g of biomass is produced per mole of ATP. In the low-energy conditions we are interested in, a single reaction may produce a quantum of energy equivalent to about one-third of a single ATP [9, 10]. If we assume that each mole of electrons from the substrate provides one mole of the energy quantum, 3.5g of biomass is produced per mole of electrons. Using the dry weight of a cell as $3 \times 10^{-13}$g [14], 3.5g dry weight is equivalent to $1.2 \times 10^{13}\text{cells} \cdot \text{mol-electron}^{-1}$. We multiply this with the flux of electron moles in the catalyzed reaction estimated above to get a rate of production of cells.

With this we can now estimate the right hand side of Eq. (S61) $\frac{\delta K_S}{h_0 E k_{cat} \mathcal{E}^{\text{ATP}} Y} \approx 4.8 \times 10^{-6}$. Thus we have $e^{\Delta G^0/RT} \ll 4.8 \times 10^{-6}$ or rather

$$-\Delta G^0 \gg 12.25RT \approx 30\text{kJ} \cdot \text{mol}^{-1}\,. \tag{S62}$$

Thus we estimate that a reaction needs to dissipating more than $30\text{kJ} \cdot \text{mol}^{-1}$ free energy for it to be effectively thermodynamically irreversible.

Due to biochemical constraints, cells assimilate at least on biological quantum of energy, about $10 - 20\text{kJ}/\,\text{mol}$ from the reaction [9, 10, 11]. Hence the overall reaction energy, which is the sum of energy dissipation and energy assimilation needs to be greater than $50\text{kJ}/\,\text{mol}$ for the reaction to be effectively irreversible. This suggests that many reactions in low-energy environments are affected by thermodynamic inhibition [17, 18, 19, 20].

Note that reactions may produce additional byproducts, such as hydrogen or methane gas [17, 18]. In such cases, improved estimates of the relevant reaction energies in the environment may be obtained by correcting for the typical concentrations of these byproducts.

### S10 An alternate prescription for relating thermodynamics and yield

A number of different studies have investigated the relation between thermodynamics of microbial metabolism to growth yield have been attempted in the literature [21, 22, 23]. Kleerebezem and Van Loosdrecht [22] have broadly classified these attempts based on their fundamental assumptions. Below we discuss two different approaches and discuss where our results apply.

In the first approach, a fixed amount of energy from catabolism is converted into ATP which is used to make biomass [21]. Production of unit biomass requires a fixed amount of ATP. Our model falls into this category, along with several others [21, 24, 15]. Since the ATP production per catabolic reaction and biomass yield per ATP is fixed, the stoichiometry relating catabolism and anabolism remains at a fixed value, which could be chosen by cell based on biochemical and stoichiometric constraints in metabolism. Due to energy assimilation into ATP in catabolism, microbial growth halts when the catabolic reaction energy drops below a nonzero threshold, consistent with experimental observations [11, 21, 25, 26].

The second approach assumes that the production of a unit mol of carbon biomass requires a fixed amount of Gibbs free energy dissipation $\Delta G_{diss}$ across all cellular processes; it uses this assumption to calculate the growth yield from catabolism [23]. Note that $\Delta G_{diss}$ represents free energy dissipated in anabolism and other processes as well, and so should not be confused with the free energy dissipation $\Delta G^0$ we use in the rest of the paper.

As explained below, this approach can be used to calculate a dynamically varying biomass yield that changes as the resource concentrations in the environment changes [27, 28]. This dynamic yield is a key element that differentiates the second approach from the first. We integrated this approach into our model and found it did not change our qualitative results (Fig. S8). Below we describe the dynamic yield approach and our implementation in greater detail.

#### The dynamic yield approach

Based on experimental observations, Heijnen and van Dijken [23] posited that the production of 1 mol C of microbial biomass required a fixed amount of Gibbs free energy dissipation $\Delta G_{diss}$ across cellular metabolic processes.The energy balance between catabolism, anabolism, and dissipation provides the following equation.

$$\lambda_{cat}\Delta G_{cat} + \Delta G_{ana} + \Delta G_{diss} = 0, \tag{S63}$$

where $\Delta G_{cat}$ is the free energy difference in the catabolic reaction with 1 mol of substrate, $\Delta G_{ana}$ is the free energy difference in the anabolic reaction that produces 1 mol C biomass, and $\lambda_{cat}$ is the stoichiometric coefficient that determines the number of catabolic reactions required for 1 mol C biomass. This equation can be rearranged to calculate the stoichiometric coefficient $\lambda_{cat}$ as

$$\lambda_{cat} = -\frac{\Delta G_{ana} + \Delta G_{diss}}{\Delta G_{cat}}. \tag{S64}$$

In the initial work [23], $\Delta G_{cat}$, $\Delta G_{ana}$ were calculated using simplified approximations to catabolism and anabolism and metabolic concentrations at the beginning of the experiment if available. Recent modeling efforts have adapted this approach to dynamically calculate $\Delta G_{cat}$ to track the change in the available catabolic free energy as the resource concentrations changes [27, 28]. This means that $\lambda_{cat}$ and hence the biomass yield per unit substrate changes dynamically. Microbial growth is then modeled using various kinetic models of catabolism coupled to this dynamic yield [27, 28].

To test how a dynamic yield affects our results, we extended our model to capture this effect. Liu et. al. [29] found that $\Delta G_{diss}$ is well-approximated by a constant value; this value is often much larger than $\Delta G_{ana}$ for biomass production from substrates such as sugars and fatty acids [27]. Motivated by these observations, we implemented the dynamic yield in our model by replacing $\Delta G_{ana} + \Delta G_{diss}$ in Eq. (S64) by a constant $A_0$. giving

$$\lambda_{cat} = -\frac{A_0}{\Delta G_{cat}}. \tag{S65}$$

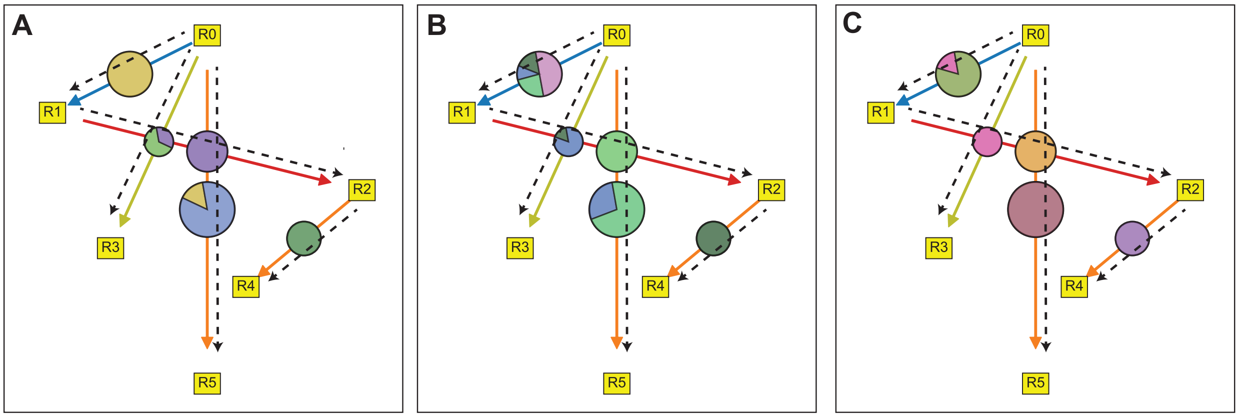


Figure S8: **Maximum dissipation principle and functional convergence in model with dynamically varying yields.** The panels show the metabolic network of three communities assembled from three different species pools in identical environments. The maximum dissipation principle predictions, shown by dashed lines, predicts the reactions accurately. The fluxes through the reactions, represented by the size of the circles on the corresponding arrows, are similar across the three communities. The pie-charts depict how the flux is shared between species in the different communities. This differs between communities. The parameters specifying the dynamical yield was $A_0/Y' = 1$. Energies of resources $R_0, R_1, R_2, R_3, R_4$, and $R_5$ were $5RT, 4RT, 3RT, 2RT, 1RT$, and $0RT$. Concentration of $R_0$ in the supply, $h_0 = 80$. The chemical kinetic parameters $K_P$,$K_S$, and $k_{cat}$ were chosen from lognormal distributions with lognormal parameters $\mu = 0, \sigma = 0.01$. Fraction of energy converted into ATP was sampled uniformly between 40% and 80%. Dilution rate, $\delta = 0.01$.

The available catabolic energy $\Delta G_{cat}$ for a reaction $P \to S$ is given by $\Delta G_{cat} = \mathcal{E}_P^0 - \mathcal{E}_S^0 + RT \ln \frac{P}{S}$. Thus as the available catabolic energy decreases, more substrate needs to be used in catabolism for the same amount of biomass production.

The dynamic yield of the second approach takes the place of the constant yield proportional to ATP production, $\mathcal{E}^{\text{ATP}} Y$ in our model equations. Therefore the microbial growth equation (Eq. (S7) ) now becomes

$$\frac{dN_\alpha}{dt} = N_\alpha \left[ \sum_{i,j \in \mathcal{X}} E_{ij}^{(\alpha)} \mathcal{F}_{ij} \frac{Y'}{\lambda_{cat}} - \delta \right], \tag{S66}$$

where $E_{ij}^{(\alpha)}$ is the enzyme allocation of species $\alpha$ to reaction $R_i \to R_j$, $\mathcal{F}_{ij}$ is the flux per enzyme in the reaction $R_i \to R_j$ defined in Eq. (S4), and $\lambda_{cat}$ is the dynamic yield from Eq. (S65), and $Y'$ is a biomass unit conversion factor. Note that while biomass production is no longer proportional to ATP production, we still need to track ATP production since it determines the free energy dissipation in catabolism which affects the flux per enzyme (Eq. (S4)).

Simulations of this model displayed functional convergence across different species pools (Fig. S8). Further, the emergent metabolic network was correctly predicted by the maximum dissipation principle. Thus our results remained qualitatively unaffected by the use of dynamic yields.

#### An illustrative example of the dynamical yield approach

The dynamical yield approach can, in principle, accommodate various details of the particular metabolic reaction being modeled. Although Including such specifics is beyond the scope our study

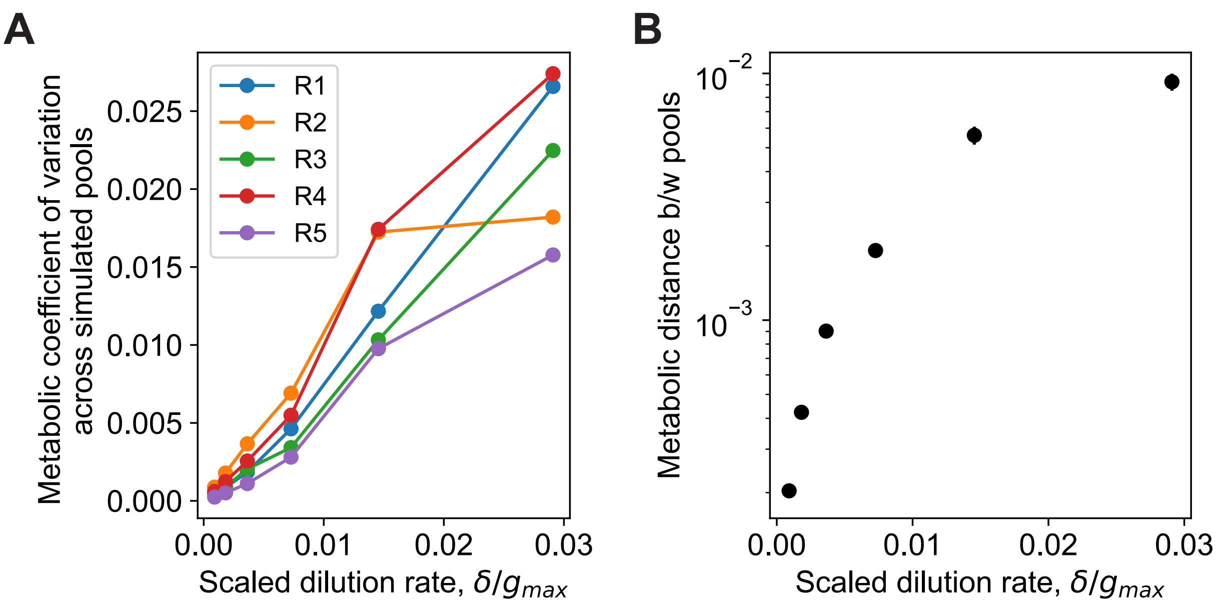


Figure S9: **Metabolic convergence weakens with dilution rate.** As seen in Fig.5 in the main text, functional convergence in both experiments and simulations weakens as dilution rate increases. Functional convergence is quantified by the coefficient of variation **(A)** and the metabolic distance **(B)**. Here simulations are run with a different set of parameters than in Fig.5. As in the experiment, the input flux of resource in simulations was held constant with $h_0\delta = 0.8$. Comparing with Fig.5, we see that the different parameters do not change the results qualitatively. Parameter here were the same as Fig.2 in the main text.

which aims to uncover general, qualitative behavior using a minimal model, we include the yield calculation for one reaction following Ref. [27] for completeness.

The reactions and energetics at pH of 7 are:

Catabolism: $1C_2H_3O_2^- + 1SO_4^{-2} \rightarrow 2HCO_3^- + 1HS^-$, with $\Delta G^{0'} = -47.7$ kJ. mol-Acetate$^{-1}$

Anabolism: $0.503C_2H_3O_2^- + 0.158NH_4^+ + 0.338\, H^+ \rightarrow 0.4305H_2O + 0.0063HCO_3^- + C_1H_{1.613}O_{0.557}\, N_{0.158}$, with $\Delta G^{0'} = 23.9$ kJ. mol-C-Biomass$^{-1}$

Dissipated energy: 432.12 kJ. mol-C-Biomass$^{-1}$. Note that this dissipated energy value is per mol of carbon biomass produced which is much larger than the energy in anabolism.

Using Eq. (S64), we get a $\lambda_{cat} = 9.6$. This would imply that from each catabolic reaction only 2.5 kJ is used in anabolism. This value is much smaller than the smallest unit of energy a cell can store, called a "quantum" of energy [9, 10, 11], which is around 10-20 kJ. Thus we conclude that at least some part of the energy assimilated from catabolism in terms of ATP is dissipated in various metabolic process. Since this dissipation is not only in catabolism, this value cannot be used as the energy dissipation Michaelis-Menten kinetics. As the concentrations of substrate and product change during the course of the experiment $\Delta G_{cat}$ can be dynamically updated [28].
